# An aptamer-mediated base editing platform for simultaneous knock-in and multiple gene knockout for allogeneic CAR-T cells generation

**DOI:** 10.1101/2023.06.20.545315

**Authors:** Immacolata Porreca, Robert Blassberg, Jennifer Harbottle, Bronwyn Joubert, Olga Mielczarek, Jesse Stombaugh, Kevin Hemphill, Jonathan Sumner, Deividas Pazeraitis, Julia Liz Touza, Margherita Francesatto, Tommaso Selmi, Juan Carlos Collantes, Zaklina Strezoska, Benjamin Taylor, Shengkan Jin, Ceri M Wiggins, Anja van Brabant Smith, John J. Lambourne

## Abstract

Gene editing technologies hold promise for enabling the next generation of adoptive cellular therapies. Conventional gene editing platforms that rely on nuclease activity, such as Clustered regularly interspaced short palindromic repeats-CRISPR associated protein 9 (CRISPR-Cas9), allow efficient introduction of genetic modifications; however, these modifications occur via the generation of DNA double-strand breaks (DSBs) and can lead to unwanted genomic alterations and genotoxicity. Here, we apply the novel modular RNA aptamer-mediated Pin-point™ base editing platform to simultaneously introduce multiple gene knockouts and site-specific integration of a transgene in human primary T cells. We demonstrate high editing efficiency and purity at all target sites and significantly reduced frequency of chromosomal translocations compared to the conventional CRISPR-Cas9 system. Site-specific knock-in of a chimeric antigen receptor (CAR) and multiplex gene knockout are achieved within a single intervention and without the requirement for additional sequence-targeting components. The ability to perform complex genome editing efficiently and precisely highlights the potential of the Pin-point platform for application in a range of advanced cell therapies.

## Introduction

Gene editing technologies have entered the clinic and show significant potential for advancing next generation therapies, particularly in the development of more efficient CAR-T cell therapies to address hematological malignancies^1–3^. To overcome the logistical and infrastructure-related challenges and product variability barriers of the autologous cell therapy paradigm, recent focus has shifted to realising the potential of allogeneic cell therapies. The manufacture of allogeneic cell products requires multiple edits to prevent both graft-versus-host disease and immune rejection by the host, which would otherwise limit efficacy and persistence of the cell product.

To expand the scope of these innovative off-the-shelf therapies to solid tumors, further edits will also be required to ensure therapeutic cells retain their efficacy in the refractory and heterogeneous tumor microenvironment^4^. These factors, together with the need to provide new functions to the cells to make effective and safe therapies that offer wider patient accessibility and therapy deployment, ultimately demand increasingly refined editing strategies.

Gene editing technologies such as zinc finger nucleases (ZFN), transcription activator-like effector nucleases (TALEN) and CRISPR-Cas9 have all been employed to successfully perform targeted editing at genomic loci for effective knockout and knock-in applications. However, the generation of DSBs inherent to their mechanism of conferring a DNA edit brings concerns of potentially deleterious mutagenic events^5–11^. The occurrence of chromosomal aberrations is enhanced in the context of multi-gene editing as more concurrent DSBs are generated, and the extent of this damage is expanded if DNA breaks also occur at off-target sites. Although many structural aberrations in a cell may not be viable, it has been reported that some rearrangements could be stable and persist over time ^1, 9^, potentially increasing the tumorigenicity risk and compromising the safety of engineered cell therapy products.

Base editing with its ability to induce genetic modifications without relying on DSB formation^12, 13^ has emerged as a strong contender in the development of advanced cell therapies, particularly in the context of multi-gene editing strategies. The two main categories of base editors, cytosine base editors (CBE) and adenine base editors (ABE) mediate efficient C to T and A to G base changes, respectively^12, 13^. Due to their capacity for programmable introduction of a single point mutation, base editors have been employed to facilitate gene disruption via generation of a premature termination codon (PTC)^14, 15^ or by mutation of splice acceptor (SA) or splice donor (SD) sites at exon-intron boundaries ^16, 17^ with high precision and efficiency whilst generating minimal undesired editing outcomes compared with standard nucleases. Rapid technological developments to increase the precision, efficiency and targeting scope of base editors, alongside an improved safety profile^16, 18, 19^, have enabled fast-tracked and successful progression to the clinic^20^. While multiple gene knockouts in immune cells have been successfully achieved by base editing ^16, 18, 19, 21^, more complex genetic modifications, such as targeted transgene integration alongside base editing knockout at other loci, have only been shown by combining two orthogonal Cas enzymes^19, 22^.

We have previously described the modular RNA aptamer-mediated Pin-point base editing system and demonstrated that this technology can edit targeted cytosines with high efficiency in human immortalized cells^23^. The Pin-point technology (Figure 1A) relies on a CRISPR-Cas module and a recruiting RNA aptamer derived from the operator stem-loop of bacteriophage MS2 (MS2) fused to the guide RNA (gRNA) to recruit the effector module. The effector module is composed of a deaminase (e.g. rAPOBEC1) fused to MS2 coat protein (MCP), which binds to the MS2 aptamer. The recruitment of the deaminase to the target site results in editing of specific residues on the unpaired DNA strand within the CRISPR R-loop.

**Figure 1.**
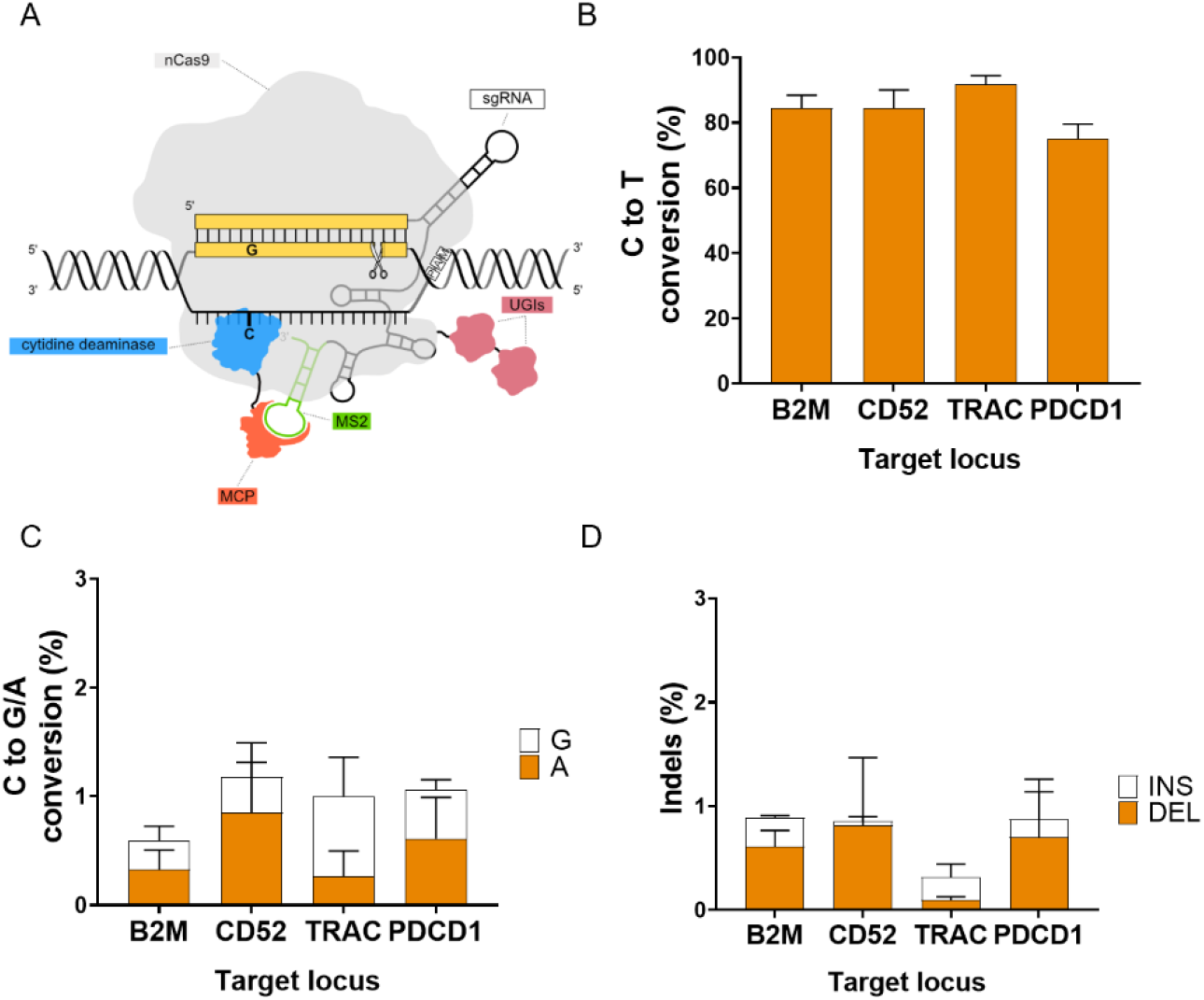
The Pin-point platform is a highly efficient technology for multiplex editing in T-cells. **A)** Schematic of the Pin-point base editing technology used in this manuscript. An SpCas9 nickase (nCas9-UGI-UGI) binds to the gRNA, the recruiting RNA aptamer (MS2) fused to the gRNA recruits the effector module. The effector module is composed of a cytidine deaminase (rAPOBEC1) fused to the aptamer binding protein (MCP). The recruitment of the deaminase to the target site forms an active complex capable of editing target cytosine residues on the unpaired DNA strand within the CRISPR R-loop. **B)** Levels of C to T conversion of the target C at *B2M, CD52, TRAC* and *PDCD1* loci following co-delivery of Pin-point mRNAs and four target sgRNAs, as analysed by NGS seven days post electroporation. **C)** Levels of C to G or A conversion of the target C at *B2M, CD52, TRAC* and *PDCD1* loci following co-delivery of Pin-point mRNAs and four target sgRNAs, as analysed by NGS. **D)** Insertion (INS) and deletion (DEL) frequency at the target C at *B2M, CD52, TRAC* and *PDCD1* loci following co-delivery of Pin-point mRNAs and four target sgRNAs, as analysed by NGS. Data represented as mean ± SD, n = 4 independent biological T-cell donors.

We demonstrate the adaptation of the plasmid-based Pin-point system^23^ into a safe and efficient fully synthetic system which can be readily adopted for manufacturing engineered cell therapies by combing mRNAs encoding the requisite Cas and deaminase modules with Pin-point gRNAs. We utilize a Pin-point base editor composed of rAPOBEC1 and Cas9 nickase (nCas9) for the generation of allogeneic human CAR-T cells. Initially, we performed a screen to identify highly functional aptamer-containing guide RNAs (gRNAs) targeting four well established genes capable of enhancing CAR-T cell function: beta-2-microglobulin (*B2M*), T cell receptor alpha constant (*TRAC*), CD52 molecule (*CD52*), and programmed cell death protein 1 (*PDCD1*). We demonstrate efficient and specific multi-gene editing with minimal differences in editing efficiencies whether editing a single locus or multiple loci and with undetectable incidence of chromosomal translocations. In addition to being compatible with conventional lentiviral CAR transgene delivery technologies the Pin-point gene editing platform can be employed to perform novel multiplex genome engineering operations, enabling simultaneous target transgene knock-in and multi targets knockout. We demonstrate the utility of this approach by combining aptamer-containing and aptamer-less gRNAs to generate functional engineered CAR-T cells via simultaneous knockout of multiple targets alongside targeted CD19-CAR insertion at the endogenous TRAC locus. The Pin-point platform thus enables complex genetic modifications of T cells using a single DNA-targeting nuclease via a novel single-step process.

## Results

### Multiplex editing in human T cells with Pin-point base editing system

To determine the optimal Pin-point system configuration with rAPOBEC1 as the effector module we assessed the impact of aptamer copy number and position within the gRNA on editing efficiencies in mammalian cells. The configuration with one copy of the MS2 aptamer located at the 3’ end of the tracrRNA resulted in optimal base editing across multiple loci (Figure S1) and was adopted as the basis for the synthetic gRNA designs employed in this study.

Using fully synthetic RNA reagents, as is conventional in engineered adoptive T cell therapy manufacturing, we screened a panel of crRNAs to identify the best performing gRNAs for knockout of *B2M*, *TRAC*, *PDCD1*, and *CD52* with the Pin-point base editing system. The crRNAs were designed to result in the introduction of a PTC or mutation at either the SA or SD sites in each target gene (Table S1). Individual crRNAs were delivered into human primary T cells by electroporation in combination with the aptamer-containing tracrRNA, an mRNA encoding nCas9 fused to a uracil glycosylase inhibitor (UGI) and an mRNA encoding rAPOBEC1-MCP. Base conversion was assessed by amplicon sequencing and target protein expression was evaluated by flow cytometry (Figure S2A-B). crRNAs exhibiting the highest level of target C to T conversion and associated protein loss for each gene (SD disruption in exon 1 for *B2M*, SD disruption in exon 1 for *CD52*, SA disruption in exon 3 for *TRAC* and SD disruption in exon 1 for *PDCD1*) were selected for synthesis as one-part single gRNAs (sgRNA) for simultaneous multi-gene editing.

Delivery of sgRNAs in multiplex achieved high levels of C to T conversion at each of the four target genes (∼76%-85%) (Figure 1B) with efficiencies comparable to that observed for individual sgRNA delivery (Figure S2C). We observed minimal undesired C to A or C to G conversion (Figure 1B-C and Figure S3A) or indel mutations (Figure 1D and Figure S3B) at each of the 4 target loci. Thus, the Pin-point system configuration consisting of nCas9 containing UGI, rAPOBEC1-MCP and an sgRNA containing an MS2 aptamer is capable of simultaneously generating C to T edits with high efficiency and purity at multiple target loci when delivered to human T cells by synthetic RNA reagents.

### Characterization of multi-gene knockout T cells

To determine the extent of multiplex target protein knockout in individual cells we performed multi color flow cytometry analysis. Consistent with the high base editing efficiency we observed at the genomic level (Figure 1B and Figure S2C), protein expression of each individual target was reduced by ∼75%-85% (Figure 2A). This was comparable to the level of protein knockout obtained using SpCas9 with sgRNAs designed for optimal indel formation at these loci (Figure 2A). Furthermore, approximately 80% of T cells edited by either the Pin-point system or by SpCas9 were negative for the three markers TCRa/b, B2M and CD52 while the remaining 20% were double negative compared to approximately 80% of mock electroporated T cells from multiple donors which were positive for all three markers (Figure 2B). PDCD1 requires T cell activation for optimal expression; therefore, T cells were cultured in the presence of phorbol 12-myristate 13-acetate (PMA) and ionomycin^24^ immediately prior to analysis to enable simultaneous quantification of all four markers. PMA-ionomycin activated T cells exhibited a more heterogeneous phenotype than unstimulated T cells due to both non-uniform upregulation of PD1 and the downregulation of TCRa/b, with ∼50% of mock electroporated controls expressing all four markers and ∼35% expressing three of the four markers (Figure S4). Nonetheless, ∼50% of T cells edited by the Pin-point system and ∼60% of T cells edited by SpCas9 were negative for the four targets, with an additional 30% negative for three of the four targets (Figure S4). This indicates a high degree of simultaneous knockout, consistent with the expectations based on individual target knockout efficiencies.

**Figure 2.**
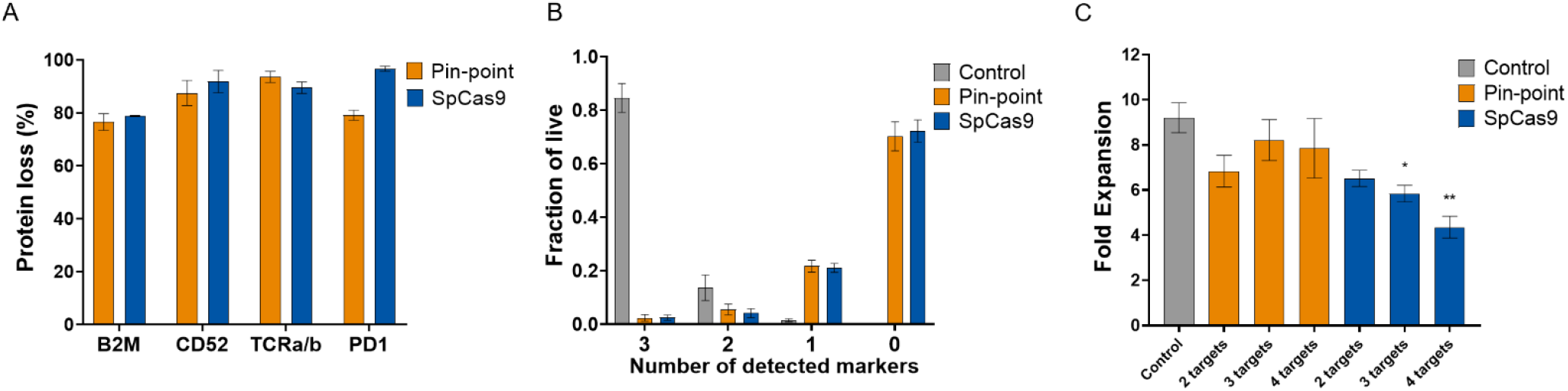
Quantification of T cell target knockout in individual cells. **A**) Frequency of CD52, TCRa/b, PD1, and B2M protein loss following co-delivery of Pin-point or SpCas9 mRNAs and their compatible four target sgRNAs, as analysed by flow cytometry seven days post electroporation. In this comparison, optimal gRNAs for SpCas9 have been used and these differ in their spacer sequence from the optimal Pin-point gRNAs (further details in the Method section). Protein loss is reported as normalised on pulse electroporated cells. **B)** Fractions of total live cells that were positive for 3 or less of three target proteins (B2M, CD52 and TCRa/b) following co-delivery of Pin-point or SpCas9 mRNAs and four target gRNAs, as analysed by flow cytometry seven days post electroporation. Control is mock electroporated T cells without RNA. **C)** Fold expansion of T cells as measured by cell counts three days post co-delivery of Pin-point or SpCas9 mRNAs and 2, 3 or 4 target gRNAs. Data represented as mean ± SD, n = 2–4 independent biological T-cell donors. *pvalue ≤0.05, **pvalue≤0.01

It is well known that nuclease-dependent gene editing technologies have the potential to impair cell fitness and proliferative capacity due to the activation of DNA-damage responses, which is exacerbated when introducing multiple DSBs^25, 26^. Consistent with the DSB-independent mechanism of base editing we observed that simultaneous editing at three or four loci with the Pin-point system did not impact T cell yield compared to SpCas9 where a significant decrease was observed (Figure 2C). Thus, the use of the Pin-point base editing technology enables efficient knockout of multiple genes in T cells without impacting cell fitness or therapeutic cell yields.

### Assessment of gRNA specific off-target editing

The potential of gene editing technologies to generate off-target edits is an important consideration for clinical risk assessment of engineered cell therapies. To experimentally identify candidate off-target editing sites for each of the four gRNAs, we performed “circularization for high-throughput analysis of nuclease genome-wide effects by sequencing” (CHANGE-seq) using SpCas9 on genomic DNA (gDNA) isolated from T cells^27^. The top 100 SpCas9 off-target candidate sites for each gRNA identified by CHANGE-seq (Table S2) were subsequently validated by rhAmpSeq in T cells edited at the four target loci using either SpCas9 or the Pin-point system. Of the 400 CHANGE-seq candidate sites that were analysed, only 2 showed detectable off-target editing by both SpCas9 and the Pin-point base editing system (1.2%-20% indel frequency and 0.7%-2.2% base editing, respectively), with an additional two sites (one for the CD52 gRNA and one for the B2M gRNA) that were edited by the Pin-point base editing system only, albeit at very low levels (0.7%-1%) (Figure 3A-B, Table S2). We therefore conclude that multi-gene editing with the Pin-point system configuration composed of nCas9 and rAPOBEC1 reduces sgRNA-dependent off-target editing compared to SpCas9.

**Figure 3.**
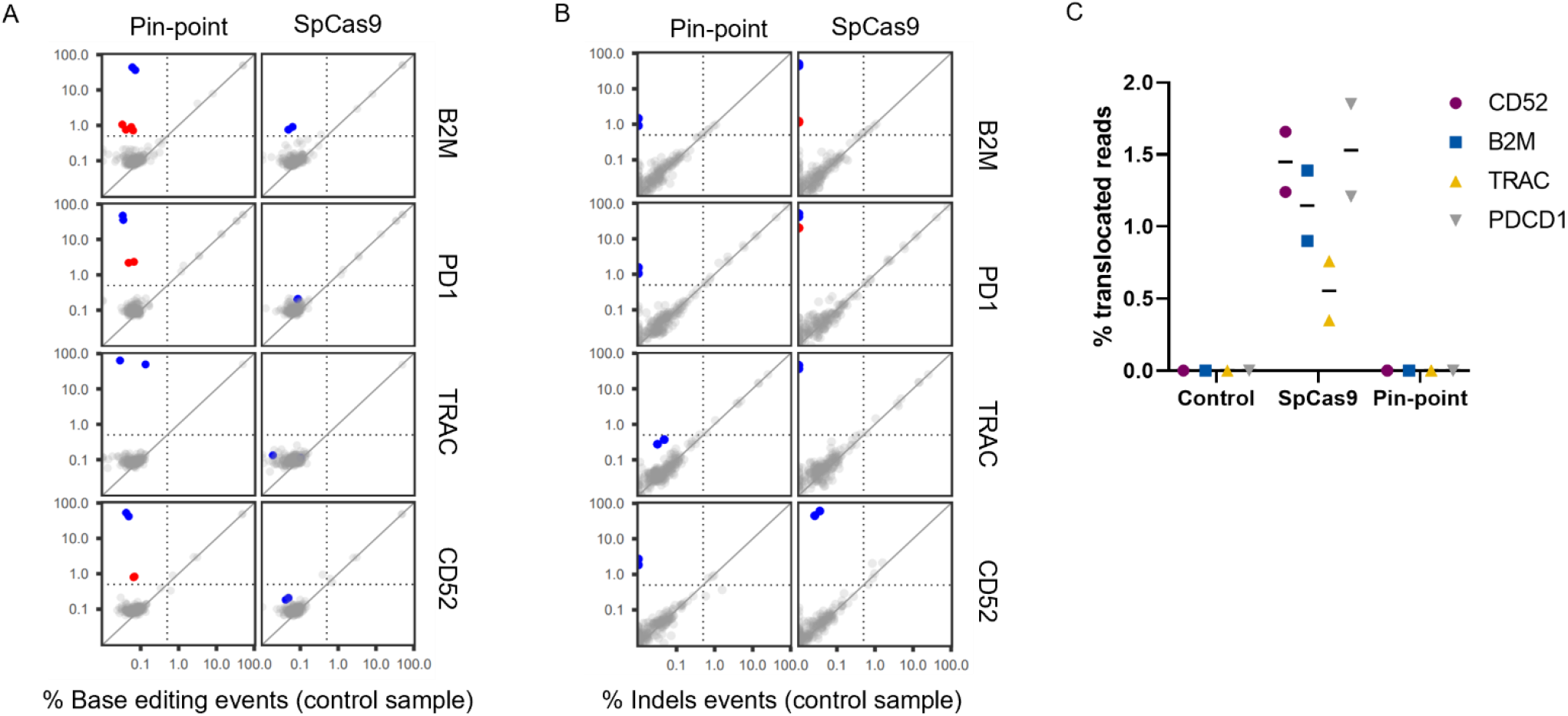
Assessment of DNA off-target editing and translocations. **A-B**) On-/off-target activity of sgRNAs targeting *B2M, PDCD1, TRAC* or *CD52* genes, determined by rhAmpSeq NGS profiling of on-target and 100 off-target sites per gRNA identified by CHANGE-seq. The on-/off-target activity of each sgRNA was profiled with either the Pin-point base editor or SpCas9 and the percent editing (% base editing events (A) or % indels events (B)) determined in each case. Each dot depicts the maximal percentage editing at a given site in one human donor for control (mock electroporation, x axis) vs edit (edited sample, y axis) with an average coverage per panel of >35,000 reads. Blue dots highlight on-target editing, while red dots highlight validated off-target activity occurring in at least 0.5 % of reads (dotted lines) and in both human donors profiled. **C)** Percentage of Capture-seq sequencing reads marked as translocations by the DRAGEN Structural Variant (SV) Caller mapping to each sgRNA target site. Pin-point or SpCas9 mRNAs were delivered with four targeting sgRNAs. Control is mock electroporated T cells without RNA. Samples were analysed three days post electroporation. n=2 independent T cell donors.

### Assessment of chromosomal translocations

In addition to the generation of undesired edits at off-target DNA sites, multiplex editing with DSB-dependent technologies can lead to the generation of chromosomal translocations ^1, 2^.

Because the Cas9 nickase variant used in the Pin-point base editing system cleaves only one DNA strand ^28, 29^, we hypothesized that multi-gene editing with the Pin-point base editor would substantially reduce the occurrence of chromosomal translocations. To test this, we performed targeted DNA capture to enrich for genomic regions around the four gRNA target sequences, followed by paired-end sequencing to an average depth of 4000X (Capture-seq) (Table S3).

Identification and quantification of translocations for each target was performed using the DRAGEN Structural Variant (SV) Caller^30 31^ (Figure 3C and Table S4). To validate the Capture-seq method, translocations between the four target genes were quantified by orthogonal droplet digital polymerase chain reaction (ddPCR) analysis using probes spanning the expected translocation breakpoints. Translocations were identified at comparable frequencies using the two methods (Figure S5A). We consistently detected all expected on-target to on-target translocation events with frequencies ranging between 0.2%-1.6%, and translocations between gRNA target regions and the PDCD1-associated off-target site identified by CHANGE-seq in the SpCas9 multi-edited samples (Figure S5B-C, Table S4), further confirming it as a contributor to CRISPR-Cas9-mediated genome instability. Frequencies of these SpCas9 induced translocations persisted over time while remaining undetectable in samples edited with the Pin-point system (Figure S5C). In summary, the aggregation of translocation frequencies quantified from T cells edited at four loci with SpCas9 (2-3%, Table S4) indicates that 1 in 33-50 haploid genomes (up to 1 in 17-25 diploid cells) will potentially carry a translocation event, while these types of chromosomal abnormalities are unlikely to occur in cells edited with the Pin-point system. Thus, multi-gene editing with the Pin-point system greatly reduces the adverse effects on genome stability associated with SpCas9.

### Molecular assessment of RNA deamination

As promiscuous activity of the deaminase component of base editors has the potential to deaminate RNA^32, 33^ similarly to the activity of endogenous cellular deaminases^34^, we assessed the impact on cytidine deamination of RNA by performing transcriptome-wide messenger RNA-sequencing (mRNA-Seq). Previous findings have highlighted that thousands of C to U transitions occur throughout the transcriptome when base editors are delivered in plasmid format^32, 33, 35^. To evaluate the impact on RNA deamination of the more therapeutically relevant RNA-based transient expression of the Pin-point system in human primary T cells, we performed an mRNA-Seq time course (day 1, 3 and 7 post-electroporation) and observed a low-level, transient, gRNA-independent increase in RNA deamination events compared to nCas9-UGI-UGI alone (approximately 60 additional C to U transitions observed exclusively at day 1 post-electroporation) (Figure 4A, B). Consistent with these observations, the level of mRNAs encoding the different components of the Pin-point base editing platform rapidly declines, becoming undetectable by day 7 in culture (Figure 4C).

**Figure 4.**
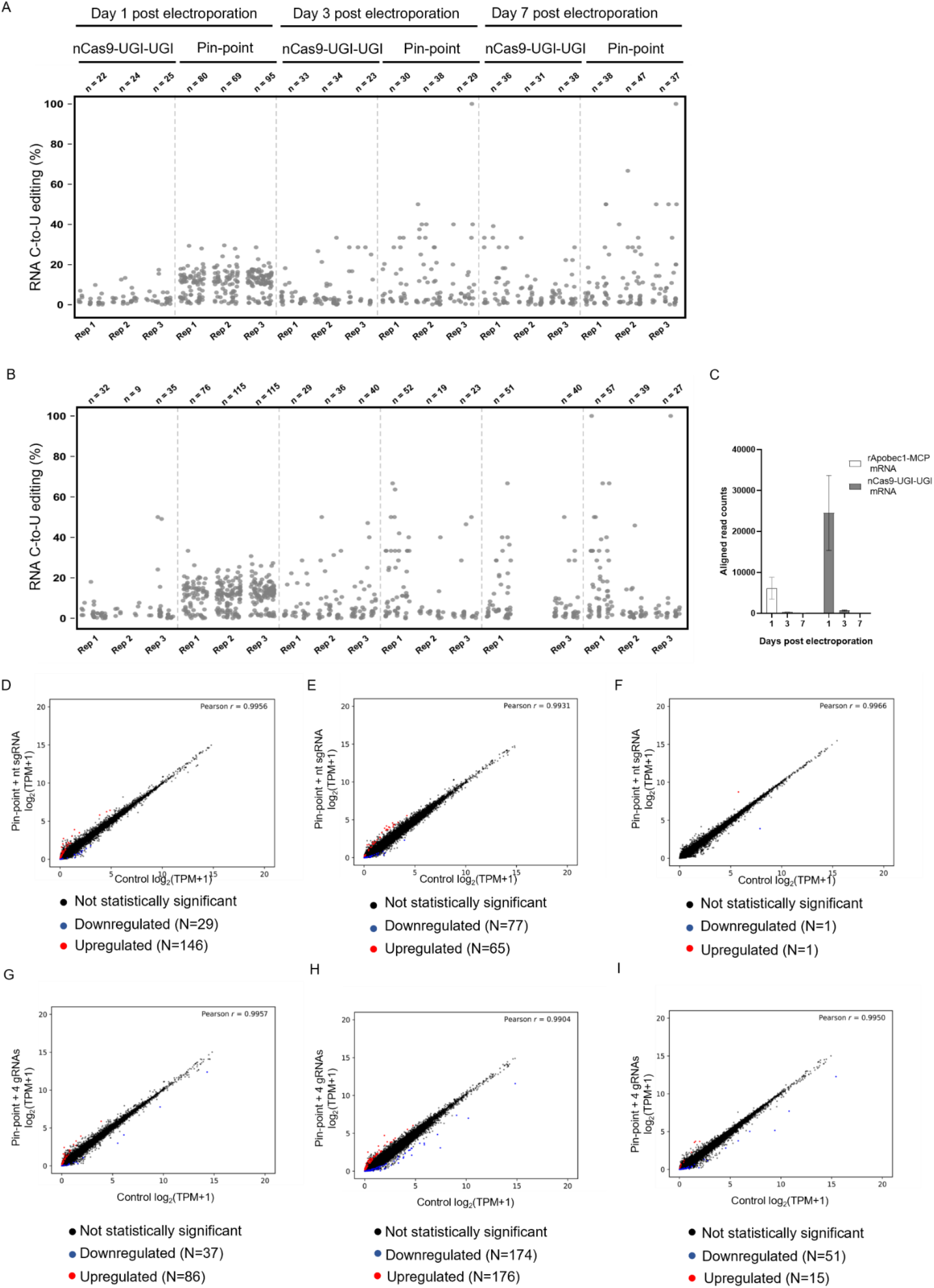
Effect of Pin-point base editing on RNA editing and transcription. **A-B**) RNA C to U editing assessed by transcriptome sequencing in primary human T cells that were electroporated with Pin-point (nCas9-UGI-UGI and rAPOBEC1-MCP) or nCas9-UGI-UGI only mRNAs and the 4 targeting sgRNAs against *B2M, CD52, TRAC* and *PDCD1* genes (A) or a scrambled non targeting sgRNA (B). Each dot represents one editing event. The total number of editing events is indicated above. **C)** Reads aligned to the Pin-point mRNA sequences (nCas9-UGI-UGI and rAPOBEC-MCP) in RNA samples from T cells electroporated with Pin-point mRNAs and the four targeting sgRNAs (*TRAC*, B2M, *CD52*, *PDCD1*) at different time points post electroporation. Individual samples were run through the GATK Best Practice for RNA-Seq Pipeline, where instead of aligning against the transcriptome, reads are aligned against the reference sequences (i.e. rAPOBEC1-MCP or nCas9-UGI-UGI) corresponding to that sample. As a result, a filtered alignment file (in BAM-format) and Variant Call Format (VCF) file was generated for each sample. Using the BAM files, read counts were determined for each component aligned against. **D-I)** Scatter plots of gene expression levels (log2 transformed TPM +1, TPM with a pseudocount of one added before log transformation) in primary human T cells electroporated with Pin-point mRNAs and either a scramble non-targeting (nt) sgRNA (D, E, F) or the four targeting sgRNAs (*TRAC*, B2M, *CD52*, *PDCD1*) (G, H,I) compared to control cells that received the pulse electroporation only (x-axis). DESeq2 analysis was performed on total mRNA collected at days 1 (D, G), 3 (E, H) and 7 (F, I) post electroporation and was used to identify up– and down-regulated genes. Up– or down-regulated genes (p < 0.05) with absolute log_2_-fold change ≥ 1.5 in gene expression (represented as log_2_ transformed TPM +1) marked red and blue, respectively. r indicates the Pearson correlation coefficient, calculated for log-transformed values on all genes.

### Phenotypic analysis of edited T cells

We investigated whether the transient mRNA deamination associated with base editing with the Pin-point system had any major effects on the gene expression profile of T cells by performing differential gene expression analysis on the mRNA-Seq time course dataset. Global gene expression was minimally affected by the delivery of Pin-point mRNAs and a non-targeting sgRNA (175, 142 and 2 transcripts were deregulated, up– or down-regulated, at day 1, 3 and 7, respectively) (Figure 4D-F and Table S5). We observed a similar effect on the transcriptome when the four gene specific sgRNAs were delivered (123, 350 and 66 transcripts were deregulated at day 1, 3 and 7, respectively) (Figure 4G-I and Table S5), indicating that the major component of the effect on gene expression profile is gRNA sequence independent.

In line with expectations five transcripts encoding the sgRNA targets B2M, PDCD1, CD52, a miRNA (MIR10393) associated with the B2M gene, and a non-coding and uncharacterized gene associated with the PDCD1 gene (LOC105373977) were stably downregulated in the samples edited with the 4 gene specific sgRNAs (Figure S6 and Table S5) whereas none of the differentially expressed transcripts were stably deregulated across the time course of samples edited with the non-targeting sgRNA. Of the deregulated genes, 41 were deregulated in both targeted and untargeted conditions one day after electroporation and 56 were deregulated in both conditions at day 3 (Figure S6). These transient gene expression changes likely reflect an immediate cellular response to the delivery of exogenous RNAs or occurred as a consequence of the transient RNA deamination events described above. In conclusion, we observed low level and transient RNA deamination by base editing using the Pin-point system that did not result in a significant long-term perturbation of the T cells transcriptional identity.

### Generation of allogeneic CAR-T cells by multi-gene editing with the Pin-point system and lentiviral delivery of the CAR

Having established that synthetic RNA-based delivery of the Pin-point base editing system presented minimal detrimental effects on human primary T cells, we employed the system to generate allogeneic CAR-T cells. We first sought to prove the compatibility of gene editing using the Pin-point system with the industry standard lentiviral CAR transgene delivery approach^36^.

Human primary T cells were first transduced with a lentivirus to deliver the CD19-CAR and then edited at the four target genes by electroporation of mRNA encoding either the Pin-point system or SpCas9 and the appropriate targeting gRNAs for *B2M*, *CD52*, *PDCD1* and *TRAC*. High efficiency protein depletion for all four targets (60-80%) was achieved, comparable to results obtained with SpCas9 (Figure 5A) without interfering with CD19-CAR expression (Figure 5B). The multi-gene edited CAR-T cells generated with the Pin-point base editing system retained the ability to kill antigen positive cancer cells in vitro (Figure 5C) and to produce the effector cytokines TNFα and IFNγ (Figure 5D) with efficiency comparable to SpCas9 edited and unedited control CAR-T cells.

**Figure 5.**
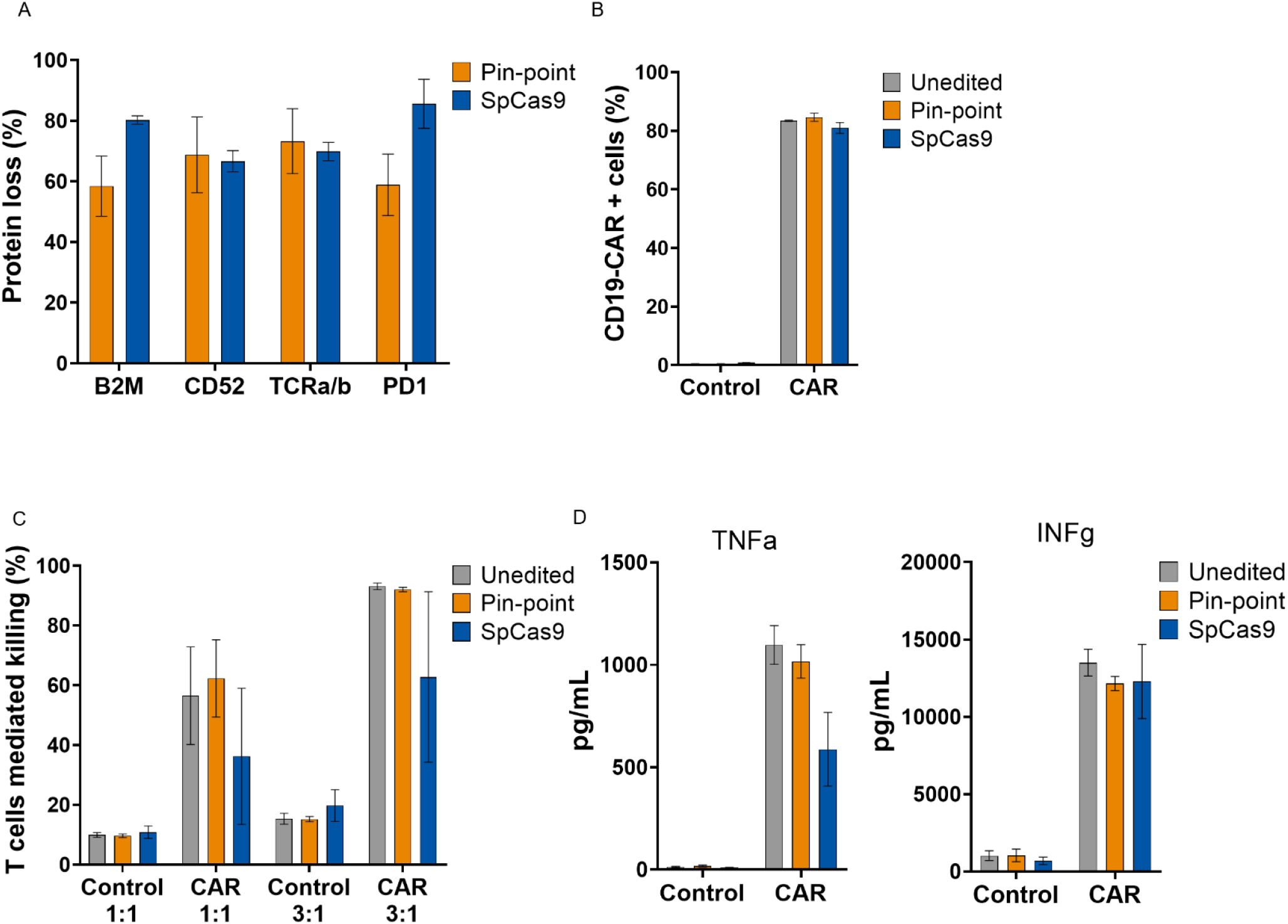
Generation of multiplex edited CAR-T cells by combining base editing by Pin-point system and lentiviral delivery of the CAR. CAR-T cells were generated by lentivirus delivery of the CD19-CAR and subsequently edited by either the Pin-point base editor or SpCas9. **A)** Frequency of CD52, TCRa/b, PD1, and B2M protein loss following co-delivery of Pin-point or SpCas9 mRNAs and four target sgRNAs, as analysed by flow cytometry seven days post electroporation in CAR-T cells. In this comparison, optimal gRNAs for SpCas9 have been used and these differ in their spacer sequence from the optimal Pin-point gRNAs (further details in the Method section). **B)** Frequency of CD19-CAR positive cells in the transduced T cell population after delivery of either Pin-point or SpCas9 reagents and in unedited cells. Control cells are T cells that have been mock transduced. **C)** Raji cells killing measured by flow cytometry after co–culture with CAR-T cells unedited or multi-edited with the Pin-point system or with SpCas9 at 1:1 or 3:1 T cells: target cells ratios. Control cells are T cells that have been mock transduced. **D)** Levels of TNFa and INFg measured in the media of the co-culture at the 1:1 T cells: target cells ratio. Data represented as mean ± SD, n = 2 independent biological T-cell donors.

### Generation of allogeneic CAR-T cells by multi-gene editing and simultaneous site-specific knock-in of the CAR with Pin-point system

In contrast to lentiviral delivery, targeted insertion of a CAR transgene can result in a more homogeneous cell therapy with improved functionality^37^ and reduced insertional oncogenesis risk. We therefore developed site-specific knock-in using the Pin-point platform by exploiting the aptamer-dependent deaminase recruitment, to achieve simultaneous multiplex gene knockout and CD19-CAR knock-in in a single event (Figure 6A). The sgRNAs containing aptamers recruit the entire Pin-point base editing machinery to the target site intended to be base edited (Figure 6A, left), while the use of two consecutive aptamer-less sgRNAs enables the recruitment of two nCas9 molecules alone at the knock-in site (Figure 6B, right) allowing to direct discreet functions to specific loci. Firstly, we verified that the deaminase expression did not affect the efficiency of site-specific knock-in of a GFP reporter or of CD19-CAR at the *TRAC* locus by the nCas9 component of the Pin-point base editing system (Figure S7A and S7B respectively).

**Figure 6.**
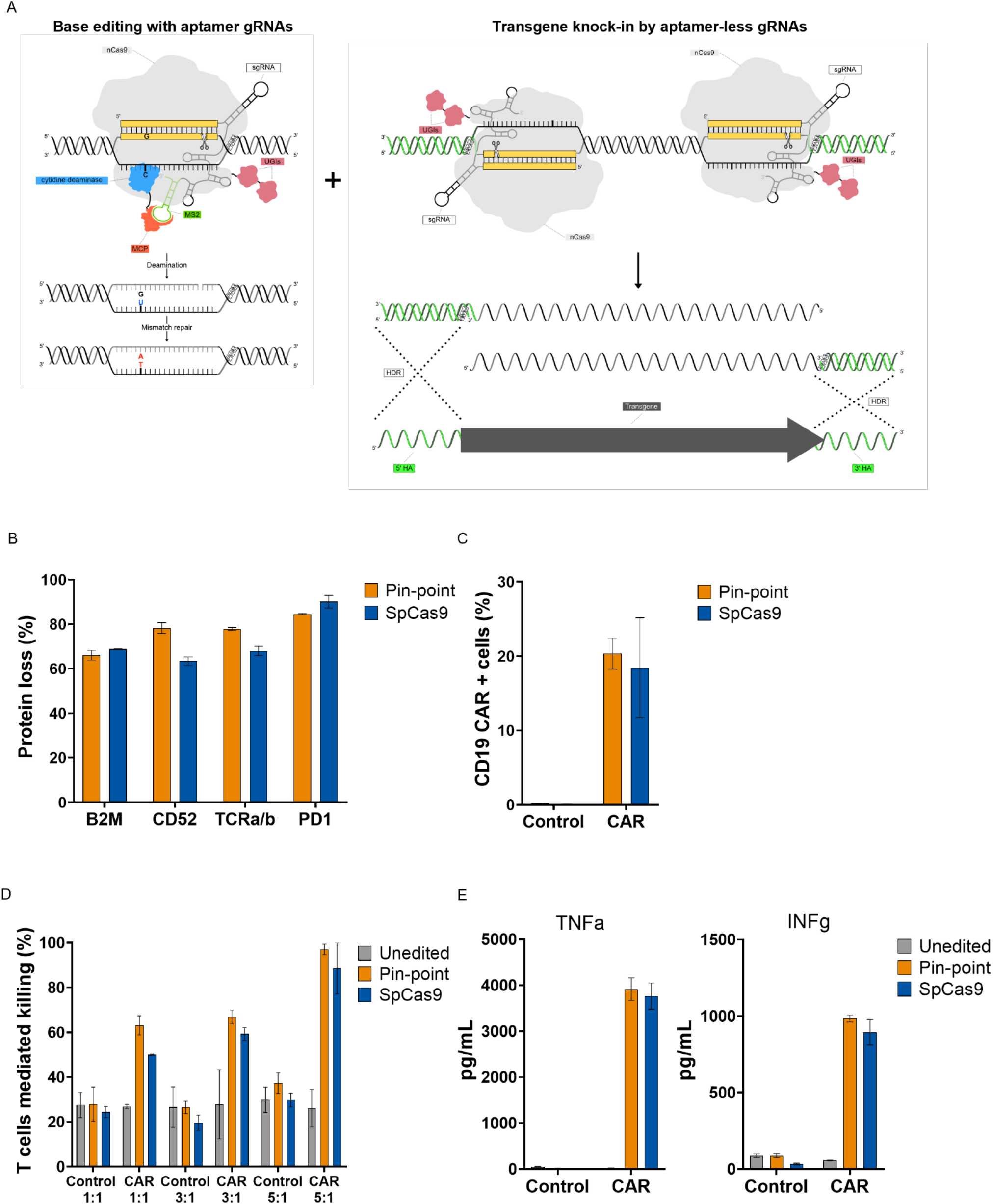
Generation of multiplex edited CAR-T cells by simultaneous multiplex base editing knockout and locus specific knock-in with the Pin-point system. **A**) Schematic showing the recruitment of the entire Pin-point system machinery by aptamer containing gRNAs on the site where the desired outcome is base editing (left) and of the nCas9 alone by aptamer-less gRNAs on the knock-in site (right). CAR-T cells were generated by knock-in of the CD19-CAR in the TRAC locus. Pin-point mRNAs have been co-delivered with aptamer containing sgRNAs directed to base edit *B2M*, *CD52* and *PDCD1* and 2 aptamer-less sgRNAs designed to target the exon1 of *TRAC* locus. Cells electroporated with SpCas9 mRNA received optimal gRNAs to knockout *B2M*, *CD52* and *PDCD1* by indels formation and one of the two gRNA designed to target the exon1 of *TRAC* locus. Shortly after electroporation, cells have been transduced with AAV6 carrying the CD19-CAR transgene flanked by the homology arms to the *TRAC* locus. **B)** Frequency of CD52, TCRa/b, PD1, and B2M protein loss following co-delivery of Pin-point or SpCas9 reagents and transduction with the AAV6-CAR as analysed by flow cytometry seven days post electroporation/transduction. **C)** Frequency of CD19-CAR positive cells in the T cell population after delivery of either Pin-point or SpCas9 reagents and transduction with the AAV6-CAR compared to non-transduced cells. **D)** Raji cells killing measured by calcein assay after co-culture with T-cells unedited or multi-edited with the Pin-point system or with SpCas9 and transduced with AAV6-CAR compared to non-transduced cells at 1:1, 3:1 or 5:1 T cells: target cells ratios. Control cells are non-transduced cells. **E)** Levels of TNFa and INFg measured in the media of the co-culture at the 1:1 T cells: target cells ratio. Data represented as mean ± SD, n = 2 independent biological T-cell donors.

Subsequently, we performed multiplex base editing using the Pin-point platform with simultaneous site-specific knock-in of CD19-CAR at the *TRAC* locus. Primary human T cells were electroporated with mRNAs encoding nCas9 and rAPOBEC1-MCP, aptamer-containing gRNAs directed to base edit *B2M*, *CD52* and *PDCD1*, and two aptamer-less gRNAs designed to target nCas9 alone to exon 1 of the *TRAC* locus to enable homology-directed repair (HDR) driven integration of the CD19-CAR. A CD19-CAR transgene lacking a promoter flanked by sequences homologous to the *TRAC* locus was then delivered by AAV6 particles. We achieved high levels (60-90%) of protein depletion for both the base editing targets (B2M, CD52 and PD1) and the integration target (TRAC) (Figure 6B). The level of site-specific knock-in evaluated by CD19-CAR expression from the endogenous TRAC locus using the simultaneous knock-in knockout application of the Pin-point system was comparable to the results achieved with SpCas9 (∼20%) (Figure 6C). Moreover, CAR-T cells generated by simultaneous knock-in and knockout using the Pin-point system were functional, showing comparable ability to kill antigen positive target cells in vitro (Figure 6D) and produce the effector cytokines TNFα and IFNγ (Figure 6E) as SpCas9 engineered controls.

These data demonstrate that the Pin-point system is a promising technology for simultaneous multiplex gene editing and targeted gene insertion applications, while limiting the deleterious effects of nuclease-dependent gene editing. Furthermore, the ability to efficiently base edit multiple sites while allowing targeted integration without the requirement of additional orthogonal targeting enzymes in a single editing procedure is unique to the Pin-point platform and has large potential in the development of complex, engineered cell and gene therapy products.

## Discussion

We present the first proof of functionality of the Pin-point system in primary human T cells, demonstrating that the technology can be employed to simultaneously introduce base edits at multiple loci at high efficiency in combination with site-specific transgene integration in a single intervention. When applied to the generation of engineered CAR-T cells using fully synthetic RNA components the Pin-point system exhibits a favourable safety profile compared to DSB-dependent CRISPR-Cas9 technology. Unbiased identification of candidate gRNA-dependent Cas9 off-target editing sites^27^ revealed the Pin-point base editing system to be highly specific with only 4 out of 400 analysed sites showing editing. When editing four targets simultaneously with SpCas9 we detected translocations at frequencies where 1 in every 17-25 cells of the final product would likely contain a translocation. Translocations were undetectable with the Pin-point technology, as it has been previously reported using other base editors ^16, 18, 19^. Multi-gene editing with the Pin-point system also improved engineered T cell yield compared to SpCas9, presenting an advantage for the manufacturing of both autologous and allogeneic therapies by increasing the yield of therapeutic product per volume of donated blood, ultimately reducing cost, opening the potential to make available at a lower price and broadening access.

Base editing combined with lentiviral delivery of the CAR transgene has been applied for multiple gene knockout for the generation of enhanced allogeneic CAR-T therapy^16, 18, 19, 21^ ^36^, however such approaches come with many limitations, including the risk of insertional mutagenesis, variable transgene expression and gene silencing. To overcome these limitations, targeted transgene integration facilitated by CRISPR-Cas technologies has become increasingly popular^38^. However, to date, simultaneous site-specific knock-in alongside base editing at other loci has only been achieved by combining two Cas homologs (i.e. Cas9 for base editing and Cas12 for knock-in) to avoid cross utilisation of sgRNAs ^19, 22^. The aptamer-dependent nature of the Pin-point system overcomes this requirement for the delivery of multiple large Cas enzymes by independently controlling which active modules are recruited at each of multiple target loci.

Whereas aptamer containing gRNAs recruit the complete base editing machinery consisting of the nCas9 and the deaminase modules to loci intended for gene knockout via base conversion, aptamer-less gRNAs recruit only the nCas9 module to loci intended for transgene insertion but avoid recruiting the deaminase function, which could otherwise induce deamination at the integration site.

Although the aptamer-dependent design of the Pin-point base editing system has the advantage of increased flexibility, the untethered deaminase could in principle increase the risk of spurious deamination. We addressed concerns about rAPOBEC1 mediated deamination^33, 35^ and determined that delivery of the Pin-point base editing machinery in the form of synthetic reagents into human primary T cells resulted in transient deamination of a minor fraction of expressed mRNAs. Nonetheless, all off-target alterations to the transcriptome rapidly dissipated and would therefore not affect the phenotype of T cells at the point of infusion of the allogeneic product. Taken together with the marked improvements in genome stability and yield of multi-gene edited T cell we propose that the Pin-point system represents a substantive advance in the toolkit available for safely engineering complex adoptive cellular therapies.

Due to its inherent flexibility, we anticipate that the Pin-point platform could be configured to simultaneously perform a suite of independent operations at multiple genomic loci by recruiting the desired effector modules via distinct RNA aptamers. For example, by combining deaminase and epigenetic modulation modules it should be possible to rewire gene regulatory networks to confer novel T cell responses to stimuli by simultaneously modifying the sequence of cis-regulatory elements and the chromatin organisation at specific loci in combination with the site-specific incorporation of synthetic signalling receptors. Similarly, it should be possible to rewire metabolic networks to overcome challenges such as T cell exhaustion by rationally engineering the activity of key enzymes in situ by base editing while simultaneously inducing or reducing expression of additional endogenous metabolic enzymes using transcriptional activator or repressor modules. Beyond its application in the creation of next-generation adoptive T cell therapies we anticipate the Pin-point system will offer similar opportunities for the engineering of a wide range of allogeneic cell therapies with increasingly advanced safety and functionality profiles.

## Material and methods

### Guide RNA design

gRNAs for base editing have been designed by using an internal design tool for PTCs generation or by manual design for the splice site disruption. The internal tool searches for NGG PAM within exons and 20bp protospacer sequences that include a C in positions 2-18 that when converted to T introduce a STOP codon. For splice site disruption, the approach was based on editing the conserved splice acceptor (intron-AG|exon) or splice donor (exon|GT-intron) motif to disrupt the functional transcript. This was done by finding an NGG PAM site near the splice junction and 20 bp protospacer that included the splice acceptor or donor site to edit. Guides that targeted more than a single location within the genome were removed from consideration. Guide RNA information for base editing is reported in Table S1. Information regarding gRNAs utilized with SpCas9 for optimal indels formation^37, 39, 40^ are reported in Table S6.

For the knock-in strategy we designed two gRNAs (Table S7) with PAM-out configuration to target opposite stands in the first exon of the TRAC gene. Both, non-homologous end joining (NHEJ) and integration of the CAR by HDR at this locus has been proven to efficiently disrupt the TCR complex^37^.

### Editing reagents

Pin-point system (nCas9-UGU-UGI and rAPOBEC1-MCP) and SpCas9 mRNAs were produced commercially (Trilink Biotechnologies and Horizon Discovery^TM^). Sequences are available in Supplemental material. Guide RNA reagents (crRNAs, tracrRNA and sgRNAs) were synthesized at Horizon^TM^, a PerkinElmer^TM^ company, or at Agilent Technologies.

### Primary human T cell isolation and culture

Primary human T cells (CD3+) were either purchased (Hemacare, CA, USA), or isolated in-house from fresh whole peripheral blood (CPD blood bags, Cambridge Bioscience, UK) or Leukopak (BioIVT) from healthy donors in accordance with Human Tissue Act (HTA) regulations. Peripheral blood mononuclear cells (PBMC) were isolated by density gradient centrifugation with Lymphoprep (StemCell Technologies, Germany) in SepMate-50 (StemCell Technologies, Germany) tubes. T cells were subsequently isolated from the PBMC population by immunomagnetic negative selective with the EasySep Human T Cell Isolation kit (StemCell Technologies, Canada). Isolated T cells with >95% viability and >95% purity were either cryopreserved or directly cultured for subsequent experiments. T cells were cultured at ∼1-2 x 10^6^mL in ImmunoCult-XF T cell expansion medium (StemCell Technologies, Canada) supplemented with Penicillin-Streptomycin (Gibco, NY, USA) and IL-2 (100 IU/mL; Miltenyi Biotech). Cells were activated with Dynabeads Human T-Activator CD3/CD28 (Gibco, Vilnius, Lithuania) at a 1:1 bead:cell ratio for 48 h prior to electroporation.

### T Cell Electroporation

After activation, Dynabeads were magnetically removed and the cells were washed with Dulbecco’s PBS (Gibco, Paisley, UK) prior to resuspension in the electroporation Buffer R. Activated T cells (2.5 x 10^5^ per reaction) were electroporated with sgRNAs at 2uM or with tracrRNA/crRNA at 6 uM and either 1 µg of SpCas9 mRNA or 1.6 µg of Pin-point nCas9-UGI-UGI and 0.2 µg of Pin-point rApobec1 using the Neon Transfection System (Invitrogen, South Korea) with the 10 µL tips and the following conditions: 1600 volts, pulse width of 10 ms, 3 pulses. After electroporation, T cells were transferred directly to prewarmed antibiotic-free ImmunoCult-XV T cell expansion medium supplemented with IL-2 (100 IU/ml), IL-7 (100 IU/ml; Peprotech, New Jersey, USA) and IL-15 (100 IU/ml; Peprotech, New Jersery, USA) and incubated at 37°C, 5% CO_2_ for 3-7 days. Electroporations were performed in duplicate or triplicate for each condition.

### Lentiviral transduction

Lentivirus was generated in HEK293T cells using Lipofectamine 3000 Transfection Reagent (Invitrogen), the ViraSafe Lentiviral Packaging System (Cell Biolabs) and an expression plasmid to deliver the 1928z CAR used in clinical trials (CD19-CAR)^41^. Viral particles were harvested from the culture, concentrated using 100 kDa Amicon® Ultra-15 Centrifugal Filter Units (Merck) and cryopreserved at –80°C. Functional viral titre was estimated by titrating the viral particles on Jurkat cells. Prior to lentiviral transduction, T cells were cultured in ImmunoCult-XV T cell expansion medium supplemented with human serum (10%; Sigma, USA), Penicillin-Streptomycin and IL-2 (100 IU/ml) and activated for 24 h in the presence of plate-bound anti-CD3 antibody (2.5 µg/mL; BioLegend) and soluble anti-CD28 antibody (2.5 µg/mL; BioLegend). Cells were transduced on RetroNectin (100 µg/mL; Takara)-coated plates at an MOI of 5 and the transduced population was enriched by puromycin (3 µg/mL; Gibco, China) selection for 5 days. Cells were then reactivated using Dynabeads Human T-Activator CD3/CD28 before electroporation with editing reagents as reported above.

### AAV transduction

The HDR donor sequence is similar to what described by Eyquem et al^37^. In more details, it consists of 1.8Kb of genomic TRAC flanking the left and the right gRNA targeting sequences, a self-cleaving P2A peptide in frame with the first exon of TRAC and by the 1928z CAR used in clinical trials^41^. The HDR sequence was cloned by GenScript in an pAAV background, and the resulting plasmid utilized to generate recombinant AAV6 donor vector by Vigene Bio.

For the locus specific knock-in experiment, activated T cells were electroporated with the gRNAs and Pin-point or SpCas9 mRNAs and immediately after electroporation transduced with the recombinant AAV6 donor vector at multiplicity of infection of 5×10^5^. Subsequently, T cells were cultured in antibiotic-free ImmunoCult-XV T cell expansion medium supplemented with IL-2 (100 IU/ml), IL-7 (100 IU/ml; Peprotech) and IL-15 (100 IU/ml; Peprotech) at 37°C, 5% CO_2_ and culture medium was completely replaced after 24 hours.

### Flow cytometry

Prior to flow cytometry, T cells edited only at the PDCD1 locus were re-stimulated using Dynabeads Human T-Activator CD3/CD28 for 48 h as described above to induce the expression of PD1. In experiments where the 4 targets were knocked-out, cells were activated with phorbol 12-myristate 13-acetate (PMA; 50 ng/mL; Sigma-Aldrich) and ionomycin (250 ng/mL; Millipore) for 48 hours prior flow cytometry analysis to induce the expression of PD1. For flow cytometry, cells were stained with fluorophore-conjugated antibodies against human B2M (BioLegend, #316304), CD52 (BD BioSciences, #562945), TCR a/b (BioLegend, #306742), PD1 (BioLegend, #329908) and CD19-CAR (AcroBiosystems, anti-FMC63 scFv). Cell viability was assessed using DAPI (80 ng/mL). Cells were acquired on an IntelliCyte IQue PLUS or Sartorius iQue3 flow cytometer using iQue ForeCyt® Enterprise Client Edition 9.0 (R3) Software for both acquisition and data analysis. The gating strategy for simultaneous quantification of viability, B2M, CD52, TCR a/b and PD1 expression was as follows (Figure S8). Within the live population, B2M expression versus CD52 expression was assessed using quadrant gating, then within each of the 4 subpopulations TCR a/b and PD1 expression was assessed using quadrant gating. Each of these 16 populations represents a different expression profile of the 4 targets and cell counts within each population were used to calculate the frequency of cells which had lost each target or combination of targets.

For fold expansion calculation, CountBright Absolute Counting Beads (Invitrogen) were added to flow cytometry samples to allow counting of the absolute number of live (DAPI negative). A flow cytometry count was performed 2h after editing (baseline), and a second one 3 days after editing. Fold expansion was calculated by dividing the live cell count for each sample by its own baseline count.

### Amplicon sequencing of genomic DNA samples

Locus-specific primers with Illumina universal adapter were designed to amplify a 250-350 bp site surrounding the genomic region of interest (Table S8). For gDNA preparation, T cells were lysed using DirectPCR (cell) (Viagen Biotech, LA, USA) lysis buffer supplemented with proteinase K (10 µg/mL; Sigma-Aldrich) and heated at 55°C for 30 min, then 95°C for 30 min. The crude lysate was then used for the first PCR with locus specific primers containing Illumina adapters. Products from the first PCR were then amplified using Illumina barcoding primers. Following barcoding, PCR samples were pooled and purified using AMPure XP beads (Beckman Coulter). DNA was sequenced by SourceBioScience on Illumina MiSeq 2 × 300 bp runs (Illumina, San Diego, CA). Illumina, San Diego, CA). Raw FASTQ files were analyzed against a reference sequence and sgRNA protospacer sequence using a custom pipeline that was used to count nucleotide substitutions in the base editor window (both expected C:G to T:A conversions and other substitutions) and indels overlapping the spacer sequence as previously described^23^.

### CHANGE-seq – Off-target discovery

CHANGE-seq was performed as previously described by Lazzarotto et al. ^27^ with minimal modifications on gDNA extracted using the Gentra Puregene Cell Kit (Qiagen) from two independent human T cell (CD3+) donors, following manufacturer’s instructions. Size analysis of resultant HMW (High Molecular Weight) gDNA was assessed in the Fragment Analyzer (Agilent),and subjected to tagmentation with customized transposome composed of oCRL225/oCRL226 adaptors and the Hyperactive Tn5 transposase (Diagenode). DNA tagmentation was performed in batches of 4ug, utilizing 17.5ul of the assembled transposome in a final volume of 200ul, and incubated for 6 minutes at 55⁰ C. Reaction was quenched by the addition of 200ul of SDS 0.4%, and resultant fragments were assessed on the Fragment analyzer and quantified by Qubit dsDNA BR Assay kit (ThermoFisher). After gap repair with Kapa Hi-Fi HotStart Uracil+ DNA Polymerase (KAPA Biosystems) and Taq DNA Ligase (NEB) and treatment with USER enzyme (NEB) and T4 polynucleotide kinase (NEB), the tagmented DNA was circularized with T4 DNA Ligase (NEB) and treated with a cocktail of exonucleases containing Plasmid-Safe ATP-dependent DNase (Lucigen), Lambda exonuclease (NEB) and Exonuclease I (NEB) to degrade residual linear DNA carryover. Circularized material was then in-vitro cleaved by SpCas9 RNP in combination with sgRNA. Illumina Universal Adaptor (NEB) was ligated to blunted end after adenylation, enzymatically treated with USER enzyme (NEB) and amplified with NEBNext Multiplex

Oligos for Illumina for 20 amplification cycles. The quality of the amplified and bead-cleaned-up libraries was determined using a 5300 fragment analyzer with the standard sensitivity NGS kit (Agilent). Libraries were then pooled, diluted, and denatured according to Illumina’s recommendations and sequenced on NextSeq550 300 cycles kit with a paired-end 2×150 configuration (Illumina). Bioinformatic analysis was performed as described by Tsai et al., 2017^42^ with a minor modification: reads with mapping quality equal to zero were included in the analysis alongside those passing the MAPQ threshold defined in the pipeline parameters, in order to nominate putative off-targets located in non-uniquely mappable regions. The pipeline was run with the following parameters: read_threshold: 4, window_size: 3, mapq_threshold: 50, start_threshold: 1, gap_threshold: 3, mismatch_threshold: 6, search_radius: 30.

### rhAMpSeq – Off-target validation

#### rhAmpSeq panel design

rhAmpSeq panels were designed for each gene target composing 100 sites with a high degree of overlap between technical and biological replicates from CHANGE-seq results. Hierarchical site selection strategy was employed to pick the most likely off-target sites: 1) sites present in both donors and all replicates, 2) sites in all replicates of one donor, 3) sites in at least two replicates of either donor, and 4) sites in at least one replicate from one donor. In cases where we had more than 100 sites we prioritized based on the nuclease-read count. The genomic coordinates for on– and off-targets were then entered into IDT’s rhAmpSeq CRISPR analysis portal for assay design and ordering.

#### rhAmpSeq library preparation

gDNA from non-edited T cells (CD3+) and T cells (CD3+) treated with either the Pin-point system or SpCas9 was extracted 5 days post electroporation using the Gentra Puregene Cell Kit (Qiagen) following manufacturer’s instructions. rhAmpSeq NGS libraries were then generated as per IDT’s rhAmpSeq library preparation protocol. Primary pools, secondary pools and single amplicon rhAmpSeq reactions were then applied on the extracted gDNA. In target rhAmp PCR 1, the 4× rhAmpSeq library mix was mixed with ∼50-80ng of gDNA and amplified using the following thermocycling conditions: 95 °C for 10 min; [95 °C for 15 s; 61 °C for 8min] × 14 cycles; 99.5 °C for 15 min; 4 °C hold. The PCR 1 product was purified using Agencourt AMPure XP beads (Beckman Coulter) and immediately proceeded to the rhAmp PCR 2. In PCR 2, dually indexed Illumina sequencing libraries were generated using PCR 1 product, mixed with 4× rhAmpSeq library mix 2 and unique i5 and i7 primers (IDT), and amplified using the following thermocycling conditions: 95°C for 3 min; [95°C for 15 s; 60°C for 30 s; 72°C for 30 s] × 24 cycles;72°C for 1 min; 4°C hold. The final libraries were purified using Agencourt AMPure XP (Beckman Coulter), quantified using Qubit 1X dsDNA HS Assay Kit (ThermoFisher Scientific) and quality was checked by qPCR and on a Tapestation 4200 (Agilent). Paired-end, 151-bp reads were sequenced using the mid-output 300 cycles kit on the Illumina’s NextSeq 550 platform (Illumina).

#### Bioinformatic processing of rhAmpSeq data

To deal with non-specific PCR products we first aligned merged reads (using FLASH: Fast Length Adjustment of SHort reads^43^) to the intended reference sequences for each target (using bwa mem) and used the alignments produced to identify the variants occurring in each reference (minimum 10 reads and allele frequency 0.01%). From these variants an extended set of reference sequences was constructed comprising the original reference plus putative variant sequences containing the different combinations of variants. From this extended set of sequences the ones which differed from the reference in global pairwise alignment by over 20 (Python Bio.pairwise2.align.globalms with scoring 1 for a match, –1 for a mismatch, 1 gap open, –0.5 gap extend) were considered sufficiently different to constitute non-specific PCR products. This set was clustered based on pairwise alignment scores within 20, and one representative from each cluster formed a “decoy” sequence to add to the targets passed to CRISPResso2 (version 2.1.1) in pooled mode with base-editing parameters (-w 20 –wc 1 –be). Following alignment, CRISPResso2 outputs were processed to identify the base position with the highest Insertion, Deletion or Base Editing event within the windows of gRNA target site +/– 10 bp per sample per amplicon. Scripts are available on request.

### Capture-seq

#### Library preparation

gDNA samples were prepared from T cells (CD3+) (Hemacare, CA, USA) using DNeasy Blood and Tissue Kit (Qiagen). 500-750ng of gDNA were used to prepare paired-end sequencing libraries using KAPA HyperPlus (Roche) workflow. Briefly, gDNA was purified using HyperPure beads (Roche), fragmented for 20mins at 37°C, and analyzed using Tapestation to confirm consistent fragmentation. Following end repair and A-tailing, KAPA universal adapters were ligated to gDNA fragments, products were cleaned and size selected using 0.8X HyperPure beads (Roche). KAPA Unique Dual Indexed primers (Roche) were incorporated by PCR (4 cycles) using KAPA HiFi Hotstart ReadyMix (Roche).

Following clean-up with HyperPure beads (Roche) libraries were quantified by Qubit and analyzed on a Tapestation 4200 (Agilent) to confirm fragment size distribution centred around 350bp.

#### Hybridisation capture probe design

DNA probes (120bp long) complementary to sequences within 200bp regions 5’ and 3’ of the PAM sites of gene editing targets were designed using ‘Oligo’ tool (https://github.com/jbkerry/oligo) in OffTarget configuration. Oligo outputs were manually curated to remove probes with stretches of homology > 30bp. 4 probes per target were chosen and positioned evenly either side of the PAM site. 5’-biotinylated DNA probes were synthesised by IDT as xGen Custom Hybridization Capture Panels.

#### Hybridisation capture & sequencing

Libraries were enriched for genomic regions flanking gene editing targets using xGen Hybridization Capture (IDT) workflow. Briefly, 1ug of each indexed library were pooled and ethanol precipitated with COT DNA, resuspended in xGen Hybridisation Buffer (IDT) containing xGen Universal Blockers TS Mix (IDT) and 5’-biotinylated custom DNA probes (IDT), heated briefly to 95°C (1 min), and hybridised overnight at 65°C. Streptavidin conjugated magnetic beads (IDT) were incubated for 45min at 65°C with the hybridised libraries, immobilised on a magnetic stand, and stringently washed at 65°C. Following washing, beads were immobilised and captured library fragments were eluted in H_2_0. Post-capture PCR (12 cycles) was performed using KAPA HiFi Hotstart ReadyMix (Roche) and xGen Library Amplification Primer Mix (IDT). Following clean-up with HyperPure beads (Roche), target enriched libraries were quantified by Qubit and analyzed by Bioanalyser to determine fragment size distribution prior to sequencing. Libraries were diluted to 1.8nM and 2×300bp paired-end reads were generated on Illumina MiSeq 2 × 300 bp runs (Illumina, San Diego) by Source Bioscience.

#### Read alignment and structural variant identification

Sequencing reads were trimmed to the first 75bps using a custom Python script and then processed through the Illumina DRAGEN Structural Variant (SV) Caller^44^ (version 3.8.4) to identify structural variants, which extends the MANTA^45^ structural variation pipeline. Each sample was run through the pipeline as an unpaired tumor sample. An example command is as follows: /opt/edico/bin/dragen –f –-ref-dir /ephemeral/ucsc.hg38.3.8.4/ –-tumor-fastq1 s3://aws_bucket/Sample1_R1_001.paired.75bp.fastq.gz –-tumor-fastq2 s3://aws_bucket/Sample1_R2_001.paired.75bp.fastq.gz –-output-directory /ephemeral/DRAGEN_Sample1/ –-output-file-prefix Sample1 –-enable-duplicate-marking true –-enable-map-align true –-enable-map-align-output true –-enable-sv true –-RGID-tumor Sample1 –-RGSM-tumor Sample1 –-sv-exome true –-remove-duplicates true

#### Translocations quantification

To remove reads derived from library fragments captured non-specifically during hybridisation the *“*.candidateSV.vcf*” output from MANTA was first filtered to include only breakends within sequences mapping to genomic regions +/– 1000bp either side of gene editing targets. Interchromosomal translocations were quantified by normalising the total count of reads (BND_PAIR_COUNT) supporting a given variant involving regions on two different chromosomes by the total number of reads mapping to either genomic region fusion point on each chromosome. Where both genomic regions adjacent to the breakpoint contained sequences targeted by capture probes the average number of reads across these regions was used for normalisation.

### Translocation quantification by ddPCR

gDNA samples were prepared from T cells (CD3+) (Hemacare, CA, USA) using DNeasy Blood and Tissue Kit (Qiagen). qPCR assays were designed to amplify predicted translocation products composed of sequences flanking each gRNA target site using the Integrated DNA Technologies (IDT) PrimeTime qPCR probe design tool. Primers and probe information for ddPCR analysis are reported in Table S9. ddPCR Supermix for Probes (no dUTP) (Bio-Rad) was used for PCR reactions each containing 40-100ng EcoR1 digested gDNA, an internal reference primer pair targeting the PPIA gene + HEX labelled probe (IDT), and a translocation targeting primer pair + FAM labelled probe (IDT). Droplets were generated and analysed using the QX200 Droplet-digital PCR system (Bio-Rad) according to manufacturer’s instructions. Translocation frequency per haploid genome was calculated from two technical replicates per sample as the fraction of translocation events detected relative to the reference sequence using QuantaSoft software (Version 1.7.4) (Bio-Rad).

### RNA purification and sequencing

Total RNA was isolated from unedited and edited T cells using the RNeasy Mini Kit (Qiagen) and quality was determined using a BioAnalyser (Agilent) and a NanoDrop Spectrophotometer.

Samples were quantified using the RNA assays on the Qubit Fluorometer. Total RNA was subjected to mRNA isolation and strand-specific RNA sequencing library preparation with the Illumina Stranded mRNA Prep, Ligation kit according to manufacturer’s instructions. The libraries were validated on the Agilent BioAnalyzer 2100 to check the size distribution of the libraries and on the Qubit to check the concentration. The RNA sequencing libraries were sequenced on an Illumina HiSeq X instrument, for an average of minimum 30M 150bp paired end reads per sample (Source Bioscience).

#### RNA deamination analysis

RNA sequence variant calling and quality control was performed as described by Grünewald et al^32^. In short, Illumina paired-end FASTQ sequences were processed through the GATK best practices for RNA-seq variant calling^46^), which produced analysis-ready BAM files aligned against human hg38 reference genome. RNA variants were called using GATK HaplotypeCaller ^47^ targeting single nucleotide variants (SNVs) across chromosomes 1-22, X and Y. Bam-readcount (https://github.com/genome/bam-readcount) was used to quantify per-base nucleotide abundances per variant.

Variant loci in the experimental samples (nCas9-UGI-UGI alone or Pin-point base editor electroporated cells) were filtered to exclude sites without high confidence reference genotype calls in the control samples. For a given SNV the read coverage in the control samples (electroporation control) was set to be above the 90th percentile of the read coverage across all SNVs in the corresponding experimental samples. Only loci having at least 99% of reads containing the reference allele in the control samples were kept. RNA edits in the experimental samples were filtered to include only loci with 10 or more reads and with greater than 0% reads containing alternate allele. Base edits labeled as C-to-U comprise C-to-U edits called on the positive strand as well as G-to-A edits sourced from the negative strand.

#### Differential gene expression analysis

Sequences were processed through the Illumina DRAGEN RNA Pipeline v3.7.5 to quantify transcripts per million and read counts. Differential expression analysis was then performed using DESeq2 v1.26.0^48^.

### Cytotoxicity assay

CD19 expressing Raji cells (InvivoGen #raji-null) were used as target cells in the cytotoxicity assay. Killing of the target cells was measured by flow cytometry assay or by calcein assay. For the flow cytometry assay, Raji cells were seeded in 96-well plate (5 x 10^4^/well) and co-cultured with T cells stained with CellTrace^TM^ Violet (Invitrogen, USA) at the indicated E:T ratios. After 3 days of coculture, cells were stained with LIVE/DEAD^TM^ Fixable Near-IR Dead Cell Stain Kit (Invitrogen, USA) and acquired by flow cytometry. With T cells positive for the CellTrace Violet, the Raji cells and T cells were gated into distinct populations prior to live-dead analysis. The percent of viable Raji cells (R_live_) was used to calculate the percent of T cell-mediated (TCM) killing as follows: TCM killing = 100 – R_live._

For the calcein assay, Raji cells were loaded with Calcein AM Dye (Invitrogen) following the manufacturer’s instructions and cocultured in 384-well plate (1 x 10^4^/well) with T cells at the indicated E:T ratios. Target cells without effectors served as a negative control and target cells incubated with 2% Triton X-100 (Merck-Sigma) served as positive control (maximum killing).

After 6 h of coculture, culture supernatant was analyzed at the Envision plate reader (PerkinElmer) with excitation 494 nm and emission 517 nm settings. TCM killing is calculated as follow: TCM killing = (test condition – negative control condition)/(positive control condition-negative control condition)*100.

### Cytokine profiling

The MultiCyt^®^ QBeads^®^ PlexScreen Secreted Protein Assay Kit (Sartorius) was used to quantify the level of tumor necrosis factor alpha (TNF-a) and interferon gamma (INF-g) secretion during the T cell cytotoxicity assays. Protocol D (reduced background, with standard curve) in the manufacturer’s handbook was followed. The IntelliCyte IQue PLUS or Sartorius iQue3 flow cytometer using iQue ForeCyt^®^ Enterprise Client Edition 9.0 (R3) Software was used for both acquisition and data analysis, including plotting the standard curves and calculating the absolute value of each sample. For samples which were diluted prior to analysis, analyte concentration was multiplied by the dilution factor. Background analyte concentration (Raji alone) was subtracted from all values and the data plotted using Graph Prism Version 9.4.1 Software.

## Acknowledgements

S.J. is supported by NJCCR Grant COCR23PRG005 and DOD Grant MD200088. We thank current Revvity employee Andrea Frapporti for creating the Pin-point system schematics used in the manuscript. Finally, we thank current and former Revvity colleagues Toby Gould, Sven Hofmann, Susana Vega, Michael Anbar, Steve Lenger, John McGonigle, Matthew Perkett for their helpful discussion and support.

## Author contributions

IP designed, performed experiments, analyzed data and wrote the manuscript. R.B., J.H., B.J., O.M. designed, performed experiments, analyzed data and contributed to the writing. K.H. designed, performed experiments and analyzed data. J. Stombaugh performed bioinformatic analysis. J. Sumner, D.P., J.L.Z., M. F. and B. T. led, designed, performed and analysed the CHANGE-Seq and rhAMpSeq – Off-target validation. Z.S., C.M.W. and A.v.B.S. supervised and provided critical suggestions. J.C.C., S.J. and T.S. contributed to the ideation of the project. J.C.C. and S.J. provided critical reagents. J.J.L. led, directed the project and contributed to the writing. R.B., J.H. and B.J. have contributed equally to the work.

