## Supplemental Material for "An aptamer-mediated base editing platform for simultaneous knock-in and multiple gene knockout for allogeneic CAR-T cells generation"

### Supplemental materials

**Table S1. Base editing guide RNA information.** crRNAs exhibiting the highest level of target C>T conversion and associated protein loss for each gene are underlined.

| Gene | spacer sequence (5'-3') | PAM | Target exon | Target C position | Base editing outcome | gRNA name |
|---|---|---|---|---|---|---|
| B2M | CACAGCCCAAGATAGTTAAG | TGG | Exon 2 | 3 | PTC | B2M_Ex2_PTC1_C3 |
| B2M | ACAGCCCAAGATAGTTAAGT | GGG | Exon 2 | 2 | PTC | B2M_Ex2_PTC2_C2 |
| B2M | TTACCCCACTTAACTATCTT | GGG | Exon 2 | 6, 7 | PTC | B2M_Ex2_PTC3_C6 |
| B2M | CTTACCCCACTTAACTATCT | TGG | Exon 2 | 7, 8 | PTC | B2M_Ex2_PTC4_C7 |
| B2M | ACTCACGCTGGATAGCCTCC | AGG | Exon 1 SD | 6 | SD disruption | B2M_Ex1_SD_C6 |
| B2M | TTGGAGTACCTGAGGAATAT | CGG | Exon 2 SA | 10 | SA disruption | B2M_Ex2_SA_C10 |
| B2M | TCGATCTATGAAAAAGACAG | TGG | Exon 3 SA | 6 | SA disruption | B2M_Ex3_SA_C6 |
| B2M | AACCTGAAAAGAAAAGAAAA | AGG | Exon 4 SA | 4 | SA disruption | B2M_Ex4_SA_C4 |
| CD52 | GTACAGGTAAGAGCAACGCC | TGG | Exon 1 | 4 | PTC | CD52_26318065_sense |
| CD52 | CTCCTCCTACAGATACAAAC | TGG | Exon 2 | 16 | PTC | CD52_26320158_sense |
| CD52 | CAGATACAAACTGGACTCTC | AGG | Exon 2 | 7 | PTC | CD52_26320167_sense |
| CD52 | CTCTTACCTGTACCATAACC | AGG | Exon 1 SD | 7 | SD disruption | CD52_Ex1_SD_anti_C7 |
| CD52 | GTATCTGTAGGAGGAGAAGT | GGG | Exon 2 SA | 5 | SA disruption | CD52_Ex2_SA_anti_C5 |
| CD52 | TGTATCTGTAGGAGGAGAAG | TGG | Exon 2 SA | 6 | SA disruption | CD52_Ex2_SA_anti_C6 |
| CD52 | GTCCAGTTTGTATCTGTAGG | AGG | Exon 2 SA | 14 | SA disruption | CD52_Ex2_SA_anti_C14 |
| TRAC | AACAAATGTGTCACAAAGTA | AGG | Exon 1 | 3 | PTC | TRAC_22547596_sense |
| TRAC | CTTCTTCCCCAGCCCAGGTA | AGG | Exon 1 | 15 | PTC | TRAC_22547761_sense |
| TRAC | TTCTTCCCCAGCCCAGGTAA | GGG | Exon 1 | 14 | PTC | TRAC_22547762_sense |
| TRAC | AGCCCAGGTAAGGGCAGCTT | TGG | Exon 1 | 5 | PTC | TRAC_22547771_sense |
| TRAC | TTTCAAAACCTGTCAGTGAT | TGG | Exon 3 | 4 | PTC | TRAC_22550570_sense |
| TRAC | TTCAAAACCTGTCAGTGATT | GGG | Exon 3 | 3 | PTC | TRAC_22550571_sense |
| TRAC | CCGAATCCTCCTCCTGAAAG | TGG | Exon 3 | 2 | PTC | TRAC_22550596_sense |
| TRAC | CTTACCTGGGCTGGGGAAGA | AGG | Exon 1 SD | 5 | SD disruption | TRAC_Ex1_SD_anti_C5 |
| TRAC | TTCGTATCTGTAAAACCAAG | AGG | Exon 3 SA | 8 | SA disruption | TRAC_Ex3_SA_anti_C8 |

| | | | | | | |
|---|---|---|---|---|---|---|
| PDCD1 | TCCAGGCATGCAGATCCCAC | AGG | Exon 1 | 11 | PTC | PDCD1_Ex1_PTC1_C11 |
| PDCD1 | TGCAGATCCCACAGGCGCCC | TGG | Exon 1 | 3 | PTC | PDCD1_Ex1_PTC2_C3 |
| PDCD1 | CGACTGGCCAGGGCGCCTGT | GGG | Exon 1 | 8, 9 | PTC | PDCD1_Ex1_PTC3_C8 |
| PDCD1 | ACGACTGGCCAGGGCGCCTG | TGG | Exon 1 | 9, 10 | PTC | PDCD1_Ex1_PTC4_C9 |
| PDCD1 | ACCGCCCAGACGACTGGCCA | GGG | Exon 1 | 6, 7 | PTC | PDCD1_Ex1_PTC5_C6 |
| PDCD1 | CACCGCCCAGACGACTGGCC | AGG | Exon 1 | 7, 8, 19, 20 | PTC | PDCD1_Ex1_PTC6_C7 |
| PDCD1 | TGTAGCACCGCCCAGACGAC | TGG | Exon 1 | 12, 13 | PTC | PDCD1_Ex1_PTC7_C12 |
| PDCD1 | GGGCGGTGCTACAACTGGGC | TGG | Exon 1 | 12 | PTC | PDCD1_Ex1_PTC8_C12 |
| PDCD1 | CGGTGCTACAACTGGGCTGG | CGG | Exon 1 | 9 | PTC | PDCD1_Ex1_PTC9_C9 |
| PDCD1 | CTACAACTGGGCTGGCGGCC | AGG | Exon 1 | 4 | PTC | PDCD1_Ex1_PTC10_C4 |
| PDCD1 | CACCTACCTAAGAACCATCC | TGG | Exon 1 | 15, 16 | PTC | PDCD1_Ex1_PTC11_C15 |
| PDCD1 | GGGGTTCCAGGGCCTGTCTG | GGG | Exon 2 | 7, 8 | PTC | PDCD1_Ex2_PTC11_C8 |
| PDCD1 | GGGGGTTCCAGGGCCTGTCT | GGG | Exon 2 | 8, 9 | PTC | PDCD1_Ex2_PTC12_C8 |
| PDCD1 | GGGGGGTTCCAGGGCCTGTC | TGG | Exon 2 | 9, 10 | PTC | PDCD1_Ex2_PTC13_C9 |
| PDCD1 | CAGCAACCAGACGGACAAGC | TGG | Exon 2 | 8 | PTC | PDCD1_Ex2_PTC14_C8 |
| PDCD1 | CCCGAGGACCGCAGCCAGCC | CGG | Exon 2 | 16 | PTC | PDCD1_Ex2_PTC15_C16 |
| PDCD1 | GGACCGCAGCCAGCCCGGCC | AGG | Exon 2 | 11 | PTC | PDCD1_Ex2_PTC16_C11 |
| PDCD1 | CGTGTCACACAACTGCCCAA | CGG | Exon 2 | 10 | PTC | PDCD1_Ex2_PTC17_C10 |
| PDCD1 | GTGTCACACAACTGCCCAAC | GGG | Exon 2 | 9 | PTC | PDCD1_Ex2_PTC18_C9 |
| PDCD1 | CGCAGATCAAAGAGAGCCTG | CGG | Exon 2 | 3 | PTC | PDCD1_Ex2_PTC19_C3 |
| PDCD1 | GCAGATCAAAGAGAGCCTGC | GGG | Exon 2 | 2 | PTC | PDCD1_Ex2_PTC20_C2 |
| PDCD1 | CACCTACCTAAGAACCATCC | TGG | Exon 1 SD | 7 | SD disruption | PDCD1_Ex1_SD_C7 |
| PDCD1 | GGAGTCTGAGAGATGGAGAG | AGG | Exon 2 SA | 6 | SA disruption | PDCD1_Ex2_SA_C6 |
| PDCD1 | TTCTTTGAGGAGAAAGGGAG | AGG | Exon 5 SA | 3 | SA disruption | PDCD1_Ex5_SA_C3 |

**Table S2. List of CHANGE-seq candidate off-targets analyzed by rhAmpSeq and editing outcomes in T cells edited with the Pin-point system or SpCas9.**

Provided as separate file

**Table S3. Sequencing depth at individual breakpoints identified by Capture-seq**

Provided as separate file

**Table S4. Frequencies of translocations identified by Capture-seq**

Provided as separate file

**Table S5. Lists of downregulated genes in T cells electroporated with Pin-point mRNAs and the 4 targets gRNAs *B2M*, *CD52*, *PDCD1* and *TRAC* or the scramble gRNA**

Provided as separate file

**Table S6. Guide RNA utilised with SpCas9 optimal for indels formation.**

| Gene | spacer sequence (5'-3') | PAM | Target exon |
|---|---|---|---|
| B2M | GAGTAGCGCGAGCACAGCTA | AGG | Exon 1 |
| CD52 | CAGCCTCCTGGTTATGGTAC | AGG | Exon 1 |
| PDCD1 | CTGCAGCTTCTCCAACACAT | CGG | Exon2 |
| TRAC | CAGGGTTCTGGATATCTGT | GGG | Exon 1 |

**Table S7. Guide RNA utilized with spCas9 for the knock-in in the TRAC locus.**

| Gene | spacer sequence (5'-3') | PAM | Target exon |
|---|---|---|---|
| TRAC | GAGAATCAAAATCGGTGAAT | AGG | Exon 1 |
| TRAC | AACAAATGTGTCACAAAGTA | AGG | Exon 1 |

**Table S8. Gene specific sequences for primers used for NGS genomic DNA amplification to detect base editing and indel events.**

| Gene | spacer sequence (5'-3') | gRNA name | Forward Primer (5'-3') | Reverse Primer (5'-3') |
|---|---|---|---|---|
| B2M | CACAGCCCAAGATAGTTAAG | B2M_Ex2_PTC1_C3 | ACTCACGTCATCCAGCAGAGA | TGGGACTCATTCAGGGTAGT |
| B2M | ACAGCCCAAGATAGTTAAGT | B2M_Ex2_PTC2_C2 | ACTCACGTCATCCAGCAGAGA | TGGGACTCATTCAGGGTAGT |
| B2M | TTACCCCACTTAACTATCTT | B2M_Ex2_PTC3_C6 | ACTCACGTCATCCAGCAGAGA | TGGGACTCATTCAGGGTAGT |
| B2M | CTTACCCCACTTAACTATCT | B2M_Ex2_PTC4_C7 | ACTCACGTCATCCAGCAGAGA | TGGGACTCATTCAGGGTAGT |
| B2M | ACTCACGCTGGATAGCCTCC | B2M_Ex1_SD_C6 | GGCCTTGTCCTGATTGGCTG | CGCTTCCCCGAGATCCAG |
| B2M | TTGGAGTACCTGAGGAATAT | B2M_Ex2_SA_C10 | AGGTGGAAGCTCATTTGGCC | ACCAGTCCTTGCTGAAAGAC |
| B2M | TCGATCTATGAAAAAGACAG | B2M_Ex3_SA_C6 | TCTGAGGCTAGTAGGAAGGGC | TCCTCAGGACAGTGAAACA |
| B2M | AACCTGAAAAGAAAAGAAAA | B2M_Ex4_SA_C4 | GGGAGCACCAAAGGGATACAC | TAAGTTGCCAGCCCTCCTA |
| CD52 | GTACAGGTAAGAGCAACGCC | CD52_26318065_sense | AAGCTGCTACCAAGACAGCC | CAGGTTTCTCTCAGGGCAGC |
| CD52 | CTCCTCCTACAGATACAAAC | CD52_26320158_sense | GAGTTCGAGACCAGCCTGAC | AGGAAAATGCCTCCGCTTAT |
| CD52 | CAGATACAAACTGGACTCTC | CD52_26320167_sense | GAGTTCGAGACCAGCCTGAC | AGGAAAATGCCTCCGCTTAT |
| CD52 | CTCTTACCTGTACCATAACC | CD52_Ex1_SD_anti_C7 | AAGCTGCTACCAAGACAGCC | CAGGTTTCTCTCAGGGCAGC |
| CD52 | GTATCTGTAGGAGGAGAAGT | CD52_Ex2_SA_anti_C5 | GAGTTCGAGACCAGCCTGAC | AGGAAAATGCCTCCGCTTAT |
| CD52 | TGTATCTGTAGGAGGAGAAG | CD52_Ex2_SA_anti_C6 | GAGTTCGAGACCAGCCTGAC | AGGAAAATGCCTCCGCTTAT |
| CD52 | GTCCAGTTTGTATCTGTAGG | CD52_Ex2_SA_anti_C14 | GAGTTCGAGACCAGCCTGAC | AGGAAAATGCCTCCGCTTAT |

| | | | | |
|---|---|---|---|---|
| TRAC | AACAAATGTGTCACAAAGTA | TRAC_22547596_sense | GCCGTGTACCAGCTGAGAGA | AAGGCCGAGACCACCAATCA |
| TRAC | CTTCTTCCCCAGCCCAGGTA | TRAC_22547761_sense | GCCGTGTACCAGCTGAGAGA | AAGGCCGAGACCACCAATCA |
| TRAC | TTCTTCCCCAGCCCAGGTAA | TRAC_22547762_sense | GCCGTGTACCAGCTGAGAGA | AAGGCCGAGACCACCAATCA |
| TRAC | AGCCCAGGTAAGGGCAGCTT | TRAC_22547771_sense | GCCGTGTACCAGCTGAGAGA | AAGGCCGAGACCACCAATCA |
| TRAC | TTTCAAAACCTGTCAGTGAT | TRAC_22550570_sense | CTGCAAGGGACAGGAGGTG | CTCACCTCAGCTGGACCAC |
| TRAC | TTCAAAACCTGTCAGTGATT | TRAC_22550571_sense | CTGCAAGGGACAGGAGGTG | CTCACCTCAGCTGGACCAC |
| TRAC | CCGAATCCTCCTCCTGAAAG | TRAC_22550596_sense | CTGCAAGGGACAGGAGGTG | CTCACCTCAGCTGGACCAC |
| TRAC | CTTACCTGGGCTGGGGAAGA | TRAC_Ex1_SD_anti_C5 | GCCGTGTACCAGCTGAGAGA | AAGGCCGAGACCACCAATCA |
| TRAC | TTCGTATCTGTAAAACCAAG | TRAC_Ex3_SA_anti_C8 | GGGGATATGCACAGAAGCTGC | CTCAGAGCTTAGGATGCACCC |
| PDCD1 | TCCAGGCATGCAGATCCCAC | PDCD1_Ex1_PTC1_C11 | CTGAGCAGTGGAGAAGGCG | CACACAGCTCAGGGTAAGGG |
| PDCD1 | TGCAGATCCCACAGGCGCCC | PDCD1_Ex1_PTC2_C3 | CTGAGCAGTGGAGAAGGCG | CACACAGCTCAGGGTAAGGG |
| PDCD1 | CGACTGGCCAGGGCGCCTGT | PDCD1_Ex1_PTC3_C8 | CTGAGCAGTGGAGAAGGCG | CACACAGCTCAGGGTAAGGG |
| PDCD1 | ACGACTGGCCAGGGCGCCTG | PDCD1_Ex1_PTC4_C9 | CTGAGCAGTGGAGAAGGCG | CACACAGCTCAGGGTAAGGG |
| PDCD1 | ACCGCCCAGACGACTGGCCA | PDCD1_Ex1_PTC5_C6 | CTGAGCAGTGGAGAAGGCG | CACACAGCTCAGGGTAAGGG |
| PDCD1 | CACCGCCCAGACGACTGGCC | PDCD1_Ex1_PTC6_C7 | CTGAGCAGTGGAGAAGGCG | CACACAGCTCAGGGTAAGGG |
| PDCD1 | TGTAGCACCGCCCAGACGAC | PDCD1_Ex1_PTC7_C12 | CTGAGCAGTGGAGAAGGCG | CACACAGCTCAGGGTAAGGG |
| PDCD1 | GGGCGGTGCTACAACTGGGC | PDCD1_Ex1_PTC8_C12 | CTGAGCAGTGGAGAAGGCG | CACACAGCTCAGGGTAAGGG |
| PDCD1 | CGGTGCTACAACTGGGCTGG | PDCD1_Ex1_PTC9_C9 | CTGAGCAGTGGAGAAGGCG | CACACAGCTCAGGGTAAGGG |
| PDCD1 | CTACAACTGGGCTGGCGGCC | PDCD1_Ex1_PTC10_C4 | CTGAGCAGTGGAGAAGGCG | CACACAGCTCAGGGTAAGGG |
| PDCD1 | CACCTACCTAAGAACCATCC | PDCD1_Ex1_PTC11_C15 | CTGAGCAGTGGAGAAGGCG | CACACAGCTCAGGGTAAGGG |
| PDCD1 | GGGGTTCCAGGGCCTGTCTG | PDCD1_Ex2_PTC11_C8 | GAAGAGGCTCTGCAGTGGAG | TGGAGAAGCTGCAGGTGAAG |
| PDCD1 | GGGGGTTCCAGGGCCTGTCT | PDCD1_Ex2_PTC12_C8 | GAAGAGGCTCTGCAGTGGAG | TGGAGAAGCTGCAGGTGAAG |
| PDCD1 | GGGGGGTTCCAGGGCCTGTC | PDCD1_Ex2_PTC13_C9 | GAAGAGGCTCTGCAGTGGAG | TGGAGAAGCTGCAGGTGAAG |
| PDCD1 | CAGCAACCAGACGGACAAGC | PDCD1_Ex2_PTC14_C8 | GGGACAACGCCACCTTCA | CAGGCTCTCTTTGATCTGCG |
| PDCD1 | CCCGAGGACCGCAGCCAGCC | PDCD1_Ex2_PTC15_C16 | GGGACAACGCCACCTTCA | CAGGCTCTCTTTGATCTGCG |
| PDCD1 | GGACCGCAGCCAGCCCGGCC | PDCD1_Ex2_PTC16_C11 | GGGACAACGCCACCTTCA | CAGGCTCTCTTTGATCTGCG |
| PDCD1 | CGTGTCACACAACTGCCCAA | PDCD1_Ex2_PTC17_C10 | GGGACAACGCCACCTTCA | CAGGCTCTCTTTGATCTGCG |
| PDCD1 | GTGTCACACAACTGCCCAAC | PDCD1_Ex2_PTC18_C9 | GGGACAACGCCACCTTCA | CAGGCTCTCTTTGATCTGCG |

| | | | | |
|---|---|---|---|---|
| PDCD1 | CGCAGATCAAAGAGAGCCTG | PDCD1_Ex2_PTC19_C3 | CAACGGGCGTGACTTCCA | GAGCTCCTGATCCTGTGCAG |
| PDCD1 | GCAGATCAAAGAGAGCCTGC | PDCD1_Ex2_PTC20_C2 | CAACGGGCGTGACTTCCA | GAGCTCCTGATCCTGTGCAG |
| PDCD1 | CACCTACCTAAGAACCATCC | PDCD1_Ex1_SD_C7 | GGCACCCTCCCTTCAACCT | CTCCAGACCCCTCGCTCC |
| PDCD1 | GGAGTCTGAGAGATGGAGAG | PDCD1_Ex2_SA_C6 | GAAGAGGCTCTGCAGTGGAG | TGGAGAAGCTGCAGGTGAAG |
| PDCD1 | TTCTTTGAGGAGAAAGGGAG | PDCD1_Ex5_SA_C3 | GAAGAGGCTCTGCAGTGGAG | TGGAGAAGCTGCAGGTGAAG |

**Table S9. Probes used for Capture-Seq**

Provided as separate file

**Table S10. Primers and probes used to detect translocation events by ddPCR**

Provided as separate file

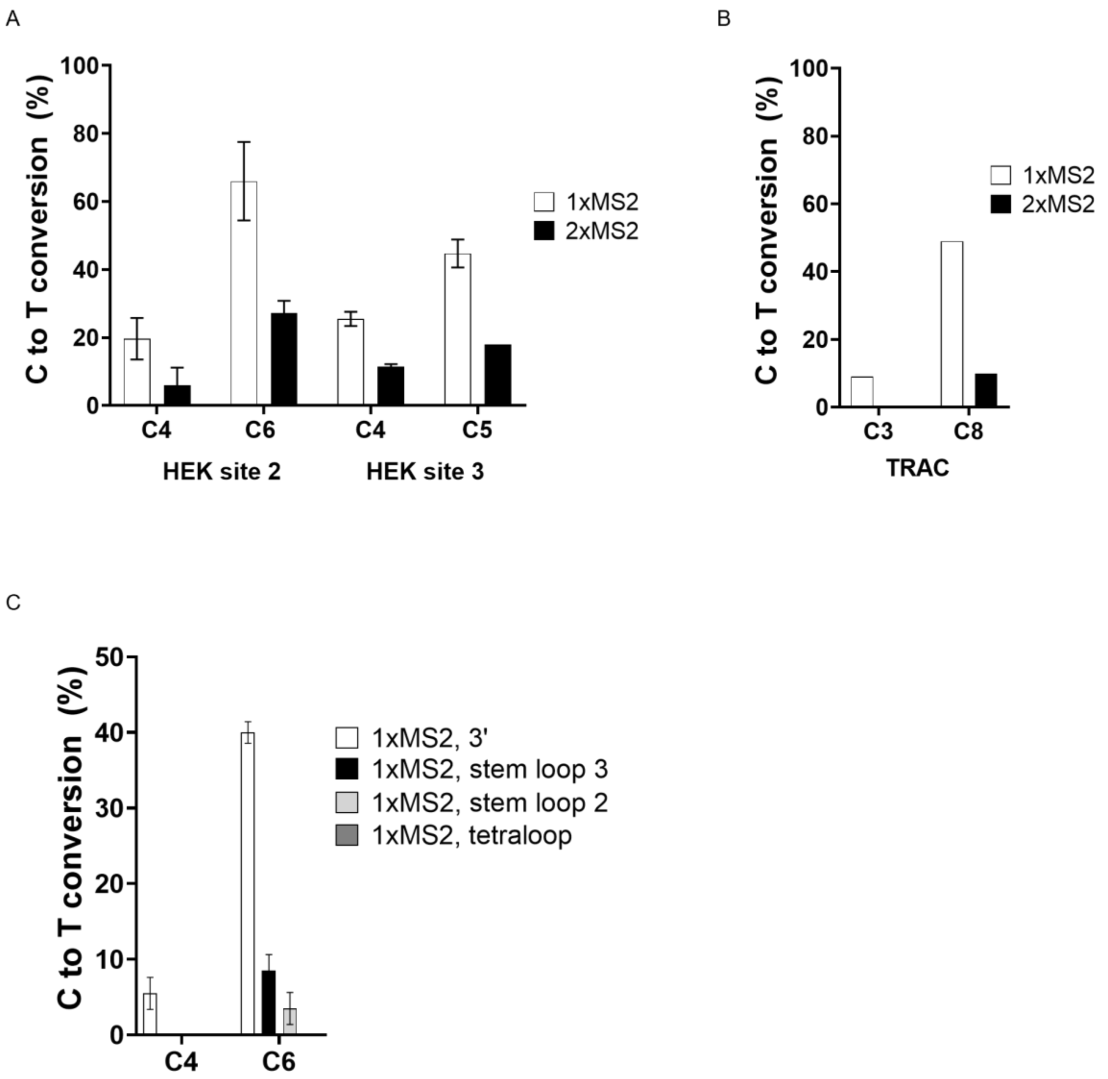


**Figure S1 Effect of one or two MS2 aptamer copies configuration on base editing activity using the Pin-point technology. A)** HEK293T cells stably expressing the nCas9 component of the Pin-point system were transfected with rAPOBEC1-MCP mRNA and the crRNA:tracrRNA complexes targeting site 2 or site 3. The tracrRNA incorporates one or two copies of the MS2 aptamer at the 3' end. Editing was analyzed by Sanger sequencing three days post-transfection. Activity for all the Cs in the editing windows is reported. **B)** Human primary T cells were electroporated with crRNA:tracrRNA complex targeting *TRAC* gene and Pin-point mRNAs. The tracrRNA incorporates one or two copies of the MS2 aptamer at the 3' end. Editing was analyzed by Sanger sequencing three days post-transfection. Preliminary results for the activity for all the Cs in the editing windows is reported. **C)** HEK293 cells were electroporated with crRNA:tracrRNA complex targeting site 2. The tracrRNA incorporates one copy of the MS2 aptamer at either the 3' terminus or at different stem-loop portions within the gRNA scaffold. Editing was analyzed by Sanger sequencing three days post-electroporation. Activity for all the Cs in the editing windows is reported.

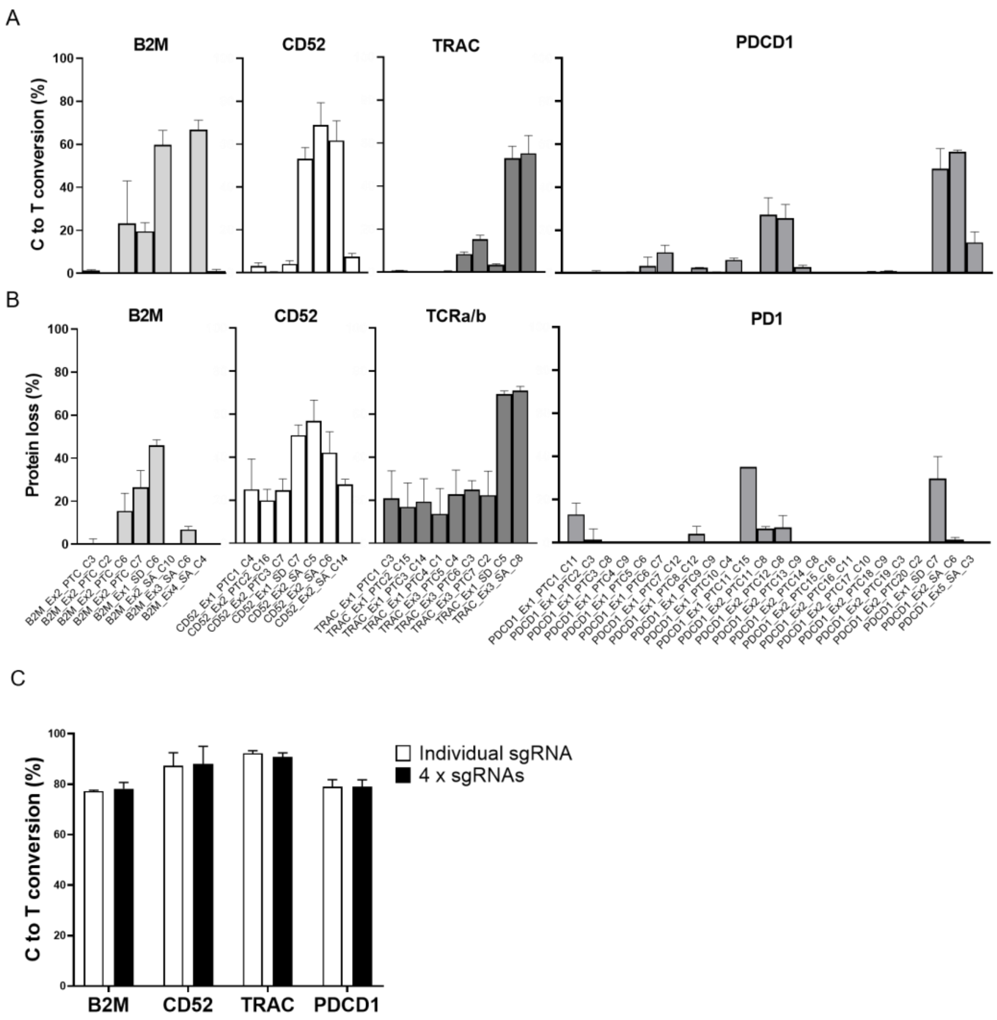


**Figure S2 Screening of optimal gRNAs for gene disruption at *CD52*, *PDCD1*, *TRAC*, and *B2M* genes by base editing using the Pin-point system. A)** Levels of C to T conversion of the target C for each of the gRNA tested at *B2M*, *CD52*, *TRAC* and *PDCD1* genes. Individual crRNAs were delivered in combination with the tracrRNA containing one copy of the MS2 aptamer at the 3’ end, nCas9-UGI-UGI mRNA, and rAPOBEC1-MCP mRNA to T cells by electroporation. Editing was analyzed by NGS three days post electroporation. **B)** Frequency of CD52, TRAC, PD1, and B2M protein loss normalised on non-electroporated samples three days after delivery of Pin-point mRNAs and individual target crRNA/tracrRNA complexes. **C)** Comparison of the levels of C to T conversion of the target C at the *B2M*, *CD52*, *TRAC* and *PDCD1* loci when optimal sgRNAs for each target are delivered individually or in combination as analyzed by Sanger sequencing seven days post electroporation. Data represented as mean ± SD, n = 2–3 independent biological T cell donors.

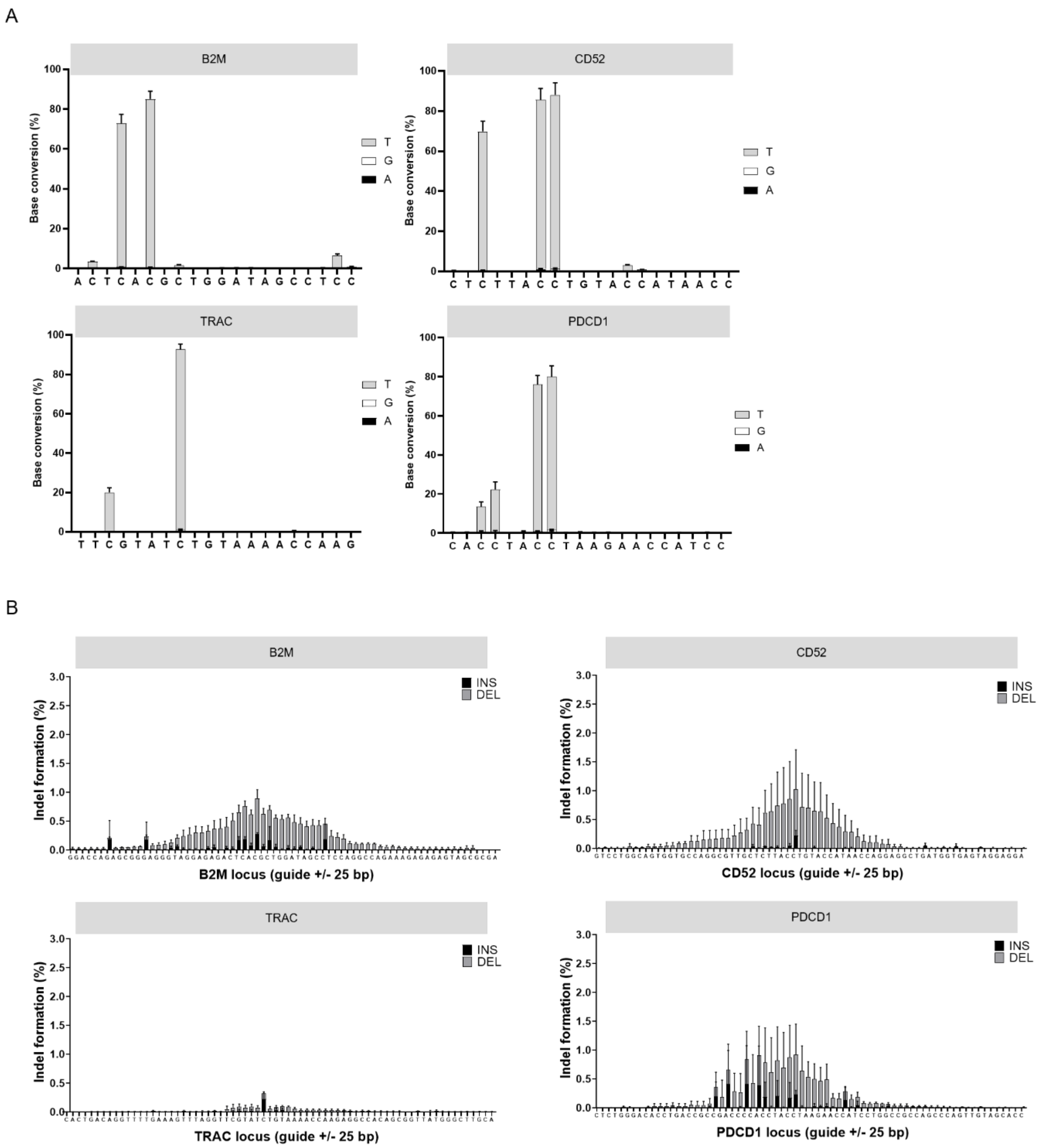


**Figure S3 Level of editing measured along the protospacer for the 4 sgRNAs. A)** Levels of base conversion (C to T, A or G) along the protospacer at *B2M*, *CD52*, *TRAC* and *PDCD1* loci following co-delivery of Pin-point mRNAs and four target sgRNAs as analyzed by NGS seven days post electroporation. **B)** Levels of insertions and deletions along the protospacer +/- 25 bp at *B2M*, *CD52*, *TRAC* and *PDCD1* loci following co-delivery of Pin-point mRNAs and four target sgRNAs as analyzed by NGS seven days post electroporation. Data represented as mean ± SD, n = 3 independent biological T cell donors.

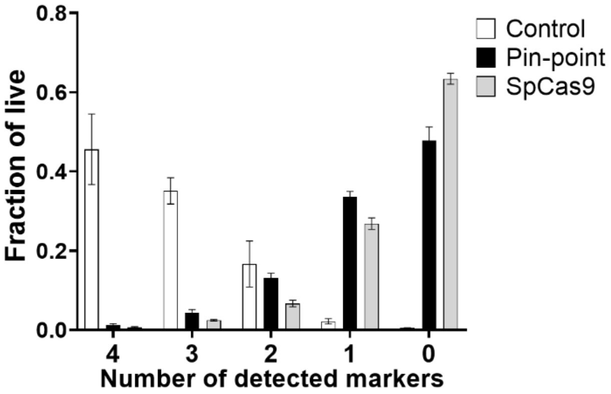


**Figure S4. Level of protein knockout for the four targets.** Fractions of cells of total live that resulted positive for 4 or less of four targets (B2M, CD52, PD1 and TCRa/b) following co-delivery of Pin-point or SpCas9 mRNAs and four target gRNAs as analyzed by flow cytometry seven days post electroporation. In this comparison, optimal sgRNAs for SpCas9 have been used and these differ in their spacer sequence from the optimal Pin-point gRNAs (further details in the Method section). To induce expression of PD1, T cells have been stimulated with PMA and ionomycin for 48 hours before flow cytometry analysis. Control is mock electroporated T cells without RNA. Data represented as mean ± SD, n = 4 independent biological T-cell donors.

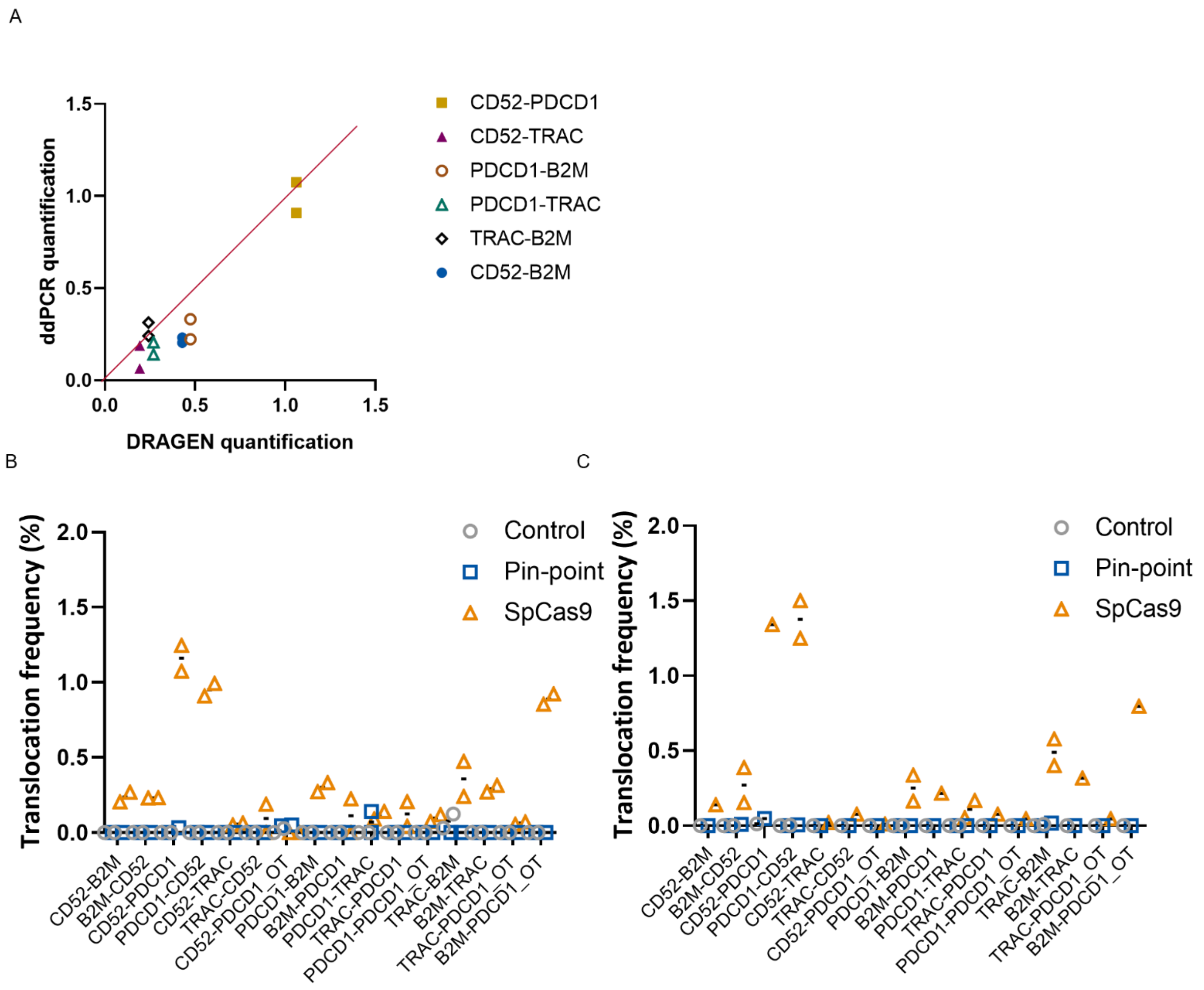


**Figure S5 ddPCR validation of translocations identified by Capture-seq. A)** Correlation of translocation frequencies between sgRNA target sites calculated from Capture-seq and ddPCR analysis of a single SpCas9 edited sample three days post electroporation. The ddPCR data points are quantifications of the two outcomes of a reciprocal translocation between two sgRNA targets. **B-C)** Individual translocation frequencies of the two outcomes of each reciprocal translocation quantified by ddPCR. Pin-point or SpCas9 encoding mRNAs were delivered with four targeting sgRNAs. Control is mock electroporated T-cells without RNA. Samples were analyzed at three (B) and seven days (C) post electroporation. n=2 independent T-cell donors.

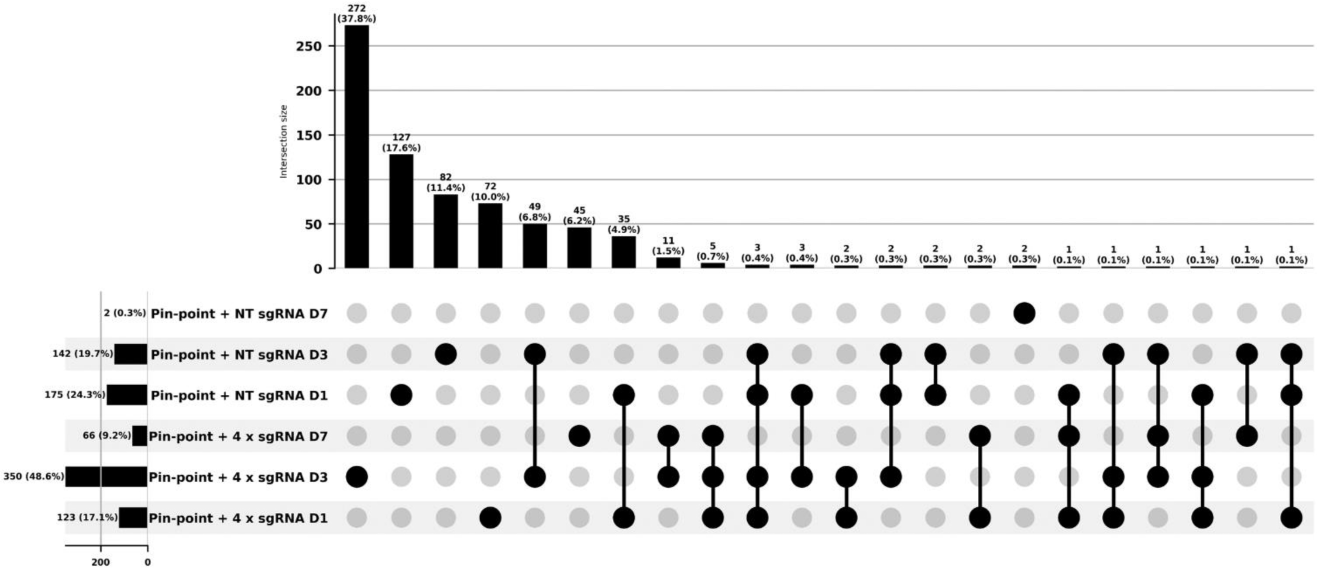


**Figure S6 UpSet plot of the differentially expressed genes (DEGs) of RNA-Seq.** This panel summarizes the deregulated genes ($p < 0.05$ and $\log_2$ FC $\geq 1.5$) overlap between human T cells electroporated with Pin-point mRNAs and either a non-targeting (NT) sgRNA or the four targeting sgRNAs and analyzed at day 1, 3 and 7 post electroporation.

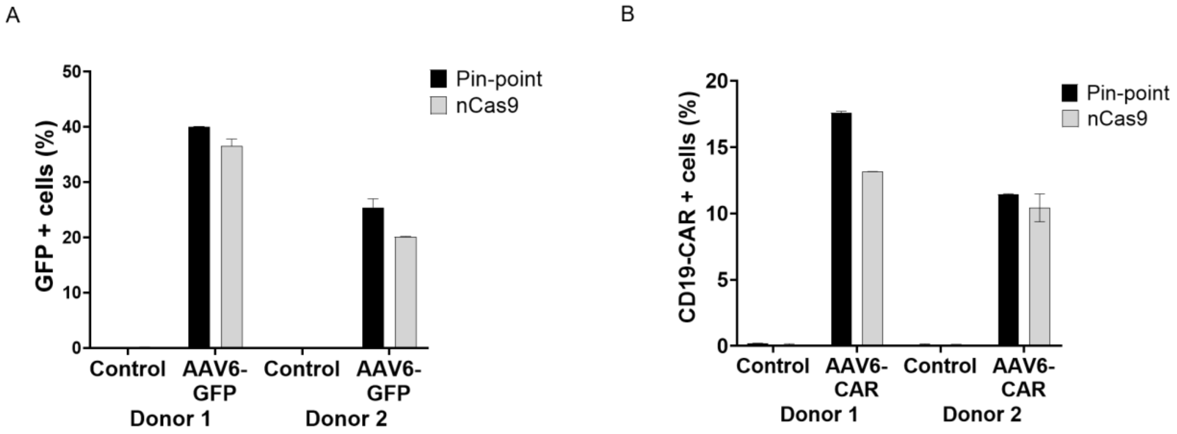


**Figure S7 Locus specific knock-in efficiency is not affected by the presence of the deaminase.** Pin-point mRNAs (nCas9-UGI-UGI and rAPOBEC1-MCP) or nCas9-UGI-UGI alone have been co-delivered with 2 aptamer-less gRNAs designed to target the exon1 of TRAC locus. Shortly after electroporation, cells have been transduced with AAV6 carrying the GFP or CD19-CAR transgene flanked by the homology arms to the TRAC locus. A) Frequency of GFP positive cells in the T cell population after delivery of either Pin-point or nCas9-UGI-UGI mRNA and transduction with the AAV6-GFP in two donors. B) Frequency of CD19-CAR positive cells in the T cell population after delivery of either Pin-point or nCas9-UGI-UGI mRNA and transduction with the AAV6-CAR in two donors.

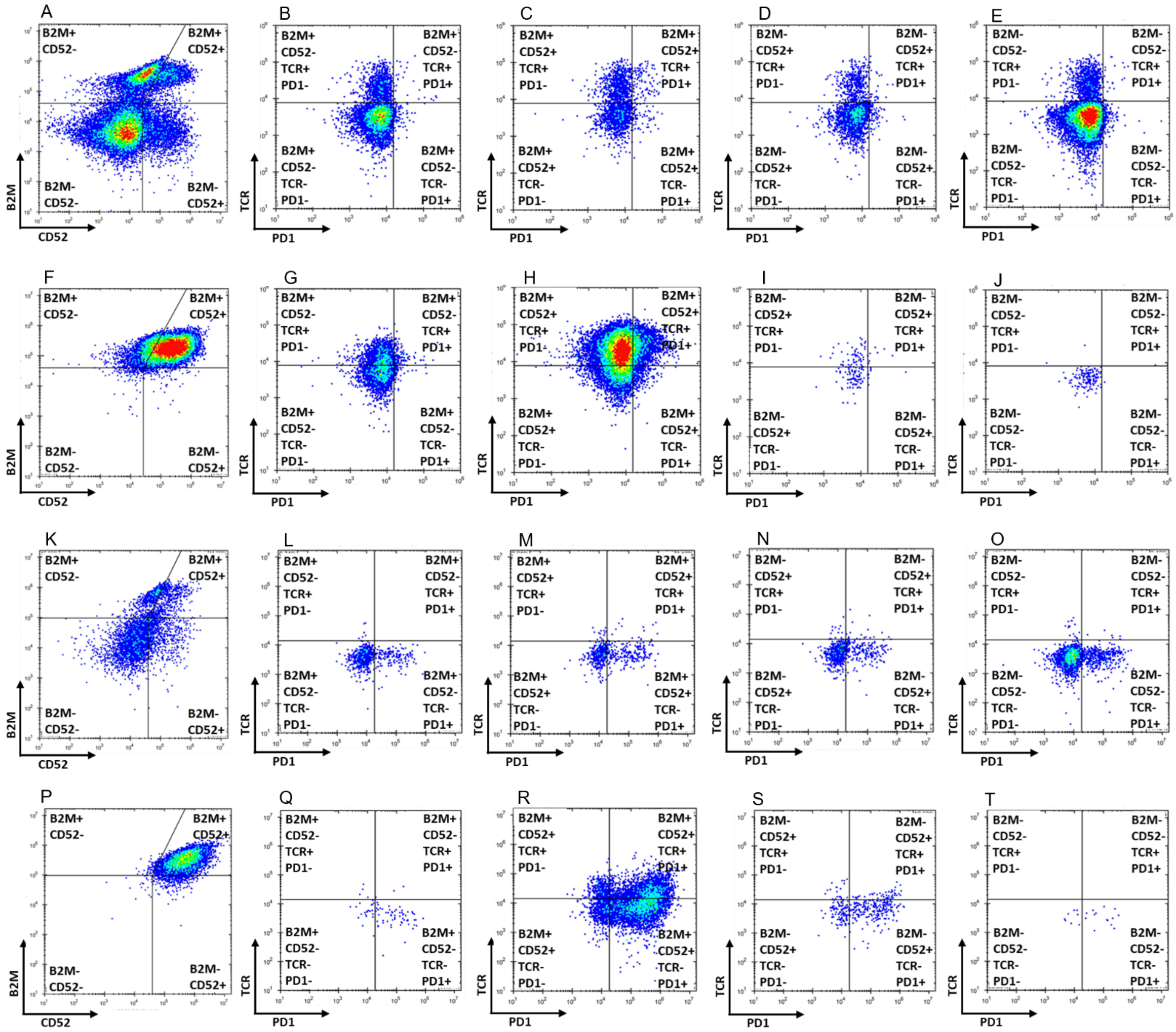


**Figure S8 Representative flow cytometry gating strategy for quadruple knockout edited and unedited cells after exclusion of doublets and dead cells.** A-E and K-O) Quadruple knockout edited cells. F-J and P to T) Unedited cells. A-J) Non stimulated cells. K-T) PMA/IO stimulated cells. A, F, K and P) B2M and CD52 expression within the live population. B, G, L and Q) TCR and PD1 expression within the B2M+ CD52- population. C, H, M and R) TCR and PD1 expression within the B2M+ CD52+ population. D, I, N and S) TCR and PD1 expression within the B2M- CD52+ population. E, J, O and T) TCR and PD1 expression within the B2M- CD52- population.

**Supplemental sequences**

**Sequence S1**. Protein sequence of $Cas9^{D10A}$

PKKKRKVDKKYSIGLAIGTNSVGWAVITDEYKVPSKKFKVLGNTDRHSIKKNLIGALLFDSGETAEATRLKRTARRR
YTRRKNRICYLQEIFSNEMAKVDDSFFHRLEESFLVEEDKKHERHPIFGNIVDEVAYHEKYPTIYHLRKKLVDSTDK
ADLRLIYLALAHMIKFRGHFLIEGDLNPDNSDVDKLFIQLVQTYNQLFEENPINASGVDAKAILSARLSKSRRLENLIA
QLPGEKKNGLFGNLIALSLGLTPNFKSNFDLAEDAKLQLSKDTYDDDLDNLLAQIGDQYADLFLAAKNLSDAILLSDI

LRVNTEITKAPLSASMIKRYDEHHQDLTLLKALVRQQLPEKYKEIFFDQSKNGYAGYIDGGASQEEFYKFIKPILEKMDGTEELLVKLNREDLLRKQRTFDNGSIPHQIHLGELHAILRRQEDFYPFLKDNREKIEKILTFRIPYYVGPLARGNSRFAWMTRKSEETITPWNFEEVVDKGASAQSFIERMTNFDKNLPNEKVLPKHSLLYEYFTVYNELTKVKYVTEGMRKPAFLSGEQKKAIVDLLFKTNRKVTVKQLKEDYFKKIECFDSVEISGVEDRFNASLGTYHDLLKIIKDKDFLDNEENEDILEDIVLTLTLFEDREMIEERLKTYAHLFDDKVMKQLKRRRYTGWGRLSRKLINGIRDKQSGKTILDFLKSDGFANRNFMQLIHDDSLTFKEDIQKAQVSGQGDSLHEHIANLAGSPAIKKGILQTVKVVDELVKVMGRHKPENIVIEMARENQTTQKGQKNSRERMKRIEEGIKELGSQILKEHPVENTQLQNEKLYLYYLQNGRDMYVDQELDINRLSDYDVDHIVPQSFLKDDSIDNKVLTRSDKNRGKSDNVPSEEVVKKMKNYWRQLLNAKLITQRKFDNLTKAERGGLSELDKAGFIKRQLVETRQITKHVAQILDSRMNTKYDENDKLIREVKVITLKSKLVSDFRKDFQFYKVREINNYHHAHDAYLNAVVGTALIKKYPKLESEFVYGDYKVYDVRKMIAKSEQEIGKATAKYFFYSNIMNFFKTEITLANGEIRKRPLIETNGETGEIVWDKGRDFATVRKVLSMPQVNIVKKTEVQTGGFSKESILPKRNSDKLIARKKDWDPKKYGGFDSPTVAYSVLVVAKVEKGKSKKLKSVKELLGITIMERSSFEKNPIDFLEAKGYKEVKKDLIIKLPKYSLFELENGRKRMLASAGELQKGNELALPSKYVNFLYLASHYEKLKGSPEDNEQKQLFVEQHKHYLDEIIEQISEFSKRVILADANLDKVLSAYNKHRDKPIREQAENIIHLFTLTNLGAPAAFKYFDTTIDRKRYTSTKEVLDATLIHQSITGLYETRIDLSQLGGDSGGSGGSGGSTNLSDIIEKETGKQLVIQESILMLPEEVEEVIGNKPESDILVHTAYDESTDENVMLLTSDAPEYKPWALVIQDSNGENKIKMLSGGSGGSGGSTNLSDIIEKETGKQLVIQESILMLPEEVEEVIGNKPESDILVHTAYDESTDENVMLLTSDAPEYKPWALVIQDSNGENKIKMLSGGSKRTADGSEFEPKKKRKV

Color key: Nuclear Localization Signal (NLS), CAS9$^{D10A}$, UGI

**Sequence S2**. Protein sequence of rAPOBEC1-MCP

PKKKRKVSSETGPVAVDPTLRRRIEPHEFEVFFDPRELRKETCLLYEINWGGRHSIWRHTSQNTNKHVEVNFIEKFTTERYFCPNTRCSITWFLSWSPCGECSRAITEFLSRYPHVTLFIYIARLYHHADPRNRQGLRDLISSGVTIQIMTEQESGYCWRNFVNYSPSNEAHWPRYPHLWVRLYVLELYCIILGLPPCLNILRRKQPQLTFFTIALQSCHYQRLPPHILWATGLKELKTPLGDTTHTSPPCPAPELLGGPMASNFTQFVLVDNGGTGDVTVAPSNFANGIAEWISSNSRSQAYKVTCSVRQSSAQNRKYTIKVEVPKGAWRSYLNMELTIPIFATNSDCELIVKAMQGLLKDGNPIPSAIAANSGIY

Color key: NLS, rApobec1, MCP

**Sequence S3.** Sequence of aptamer containing tracrRNA

AACAGCAUAGCAAGUUAAAAUAAGGCUAGUCCGUUAUCAACUUGAAAAAGUGGCACCGAGUCGGUGCGCGCACAUGAGGAUCACCCAUGUGCUUUUmU*mU*U

Color key: gRNA scaffold, MS2 aptamer, mN* nucleotides containing 2′-O-methyl 3′phosphorothioate modifications.

**Sequence S4.** Sequence of aptamer containing sgRNA

mN*mN*NNNNNNNNNNNNNNNNNNGUUUUAGAGCUAGAAAUAGCAAGUUAAAAUAAGGCUAGUCCGUUAUCAACUUGAAAAAGUGGCACCGAGUCGGUGCGCGCACAUGAGGAUCACCCAUGUGCUUUUmU*mU*U

Color key: Spacer sequence, gRNA scaffold, MS2 aptamer, mN* nucleotides containing 2′-O-methyl 3′phosphorothioate modifications.

**Sequence S5.** Sequence of aptamer-less sgRNA

mN*mN*NNNNNNNNNNNNNNNNNNGUUUUAGAGCUAGAAAUAGCAAGUUAAAAUAAGGCUAGUCCGUUAUCAACUUGAAAAAGUGGCACCGAGUCGGUGCUmU*mU*U

Color key: Spacer sequence, gRNA scaffold, mN* nucleotides containing 2′-O-methyl 3′phosphorothioate modifications.
